# Free energy landscapes of KcsA inactivation

**DOI:** 10.1101/2023.04.05.535698

**Authors:** Sergio Pérez-Conesa, Lucie Delemotte

## Abstract

The bacterial ion channel KcsA has become a useful model of complex K+-ion channels thanks to its single pore domain structure whose sequence shares many similarities with eukaryotic channels. Like many physiologically-relevant ion channels, KcsA inactivates after prolonged exposure to stimuli (in this case, a lowered pH). The inactivation mechanism has been heavily investigated, using structural, functional and simulations methods, but the molecular basis underlying the energetics of the process remains actively debated. In this work, we use the “string method with swarms of trajectories” enhanced sampling technique to characterize the free energy landscape lining the KcsA inactivation process. After channel opening following a pH drop, KcsA presents metastable open states leading to an inactivated state. The final inactivation step consists of a constriction of the selectivty filter and entry of three water molecules into binding sites behind each selectivity filter subunit. Based our simulations, we propose a key role for residue L81 in opening a gateway for water molecules to enter their buried sites, rather than for Y82 which has previously been suggested to act as a lid. In addition, since we found the energetically favored inactivation mechanism to be dependent on the force field, our results also address the importance of parameter choice for this type of mechanism. In particular, inactivation involves passing through the fully-open state only when using the AMBER force field. In contrast, using CHARMM, selectivity filter constriction proceeds directly from the partially open state. Finally, our simulations suggest that removing the co-purifying lipids stabilizes the partially open states, rationalizing their importance for the proper inactivation of the channel.

## Introduction

The bacterial ion channel KcsA has become a standard model for complex eukaryotic K-ion channels due to the highly conserved nature of its sequence and the relative simplicity of its structure consisting of a single pore domain (***Figure 1a-b***). Indeed, the characteristic TVGYG sequence of the selectivity filter plays an important role in determining channel selectivity(***Doyle et al., 1998***). When the homotetramer is formed, the selectivity filter has four potassium binding sites (S1 to S4) which select for K+ over Na+ with a remarkable selectivity (***Figure 1a***). KcsA features two main regions where the flow of ions can be interrupted: the helical bundle crossing of TM2 or inner gate (IG) and the selectivity filter (SF), which can stop ion flow through its constriction at the level of residue G77(***Li et al., 2018***). One of the hallmarks of ion channel biochemistry is channel inactivation after prolonged exposure to the opening stimulus (***Hille, 2001***). This mechanism is essential to many physiological signaling mechanisms. In KcsA, the inactivation mechanism appears fairly established (***Li et al., 2018***), largely on the basis of numerous X-Ray diffraction structures (***Figure 1a-d***) (***Doyle et al., 1998; Cuello et al., 2010a,b, 2017***). Upon exposure to low pH, the inner gate opens, allowing current to flow through the channel. The protein then visits millisecond-lived metastable states of increasing degrees of inner gate opening (***Figure 1b***), until an allosteric signal is transmitted from the inner gate to the selectivity filter causing it to constrict, ceasing the current and forming the inactivated state (***Figure 1c***). In this way, the selectivity filter acts as a secondary gate which is allosterically connected to the inner gate. In the conductive state, a single water molecule is bound behind each subunit of the selectivity filter. The final inactivation step is accompanied by the binding of two additional water molecules, leading to a characteristic configuration with three water molecules bound behind the constricted selectivity filter (***Ostmeyer et al., 2013***). These waters have been described to go through a gate formed by the Y82 “lid” residue (***Ostmeyer et al., 2013***; ***Cuello et al., 2017***).

**Figure 1.**
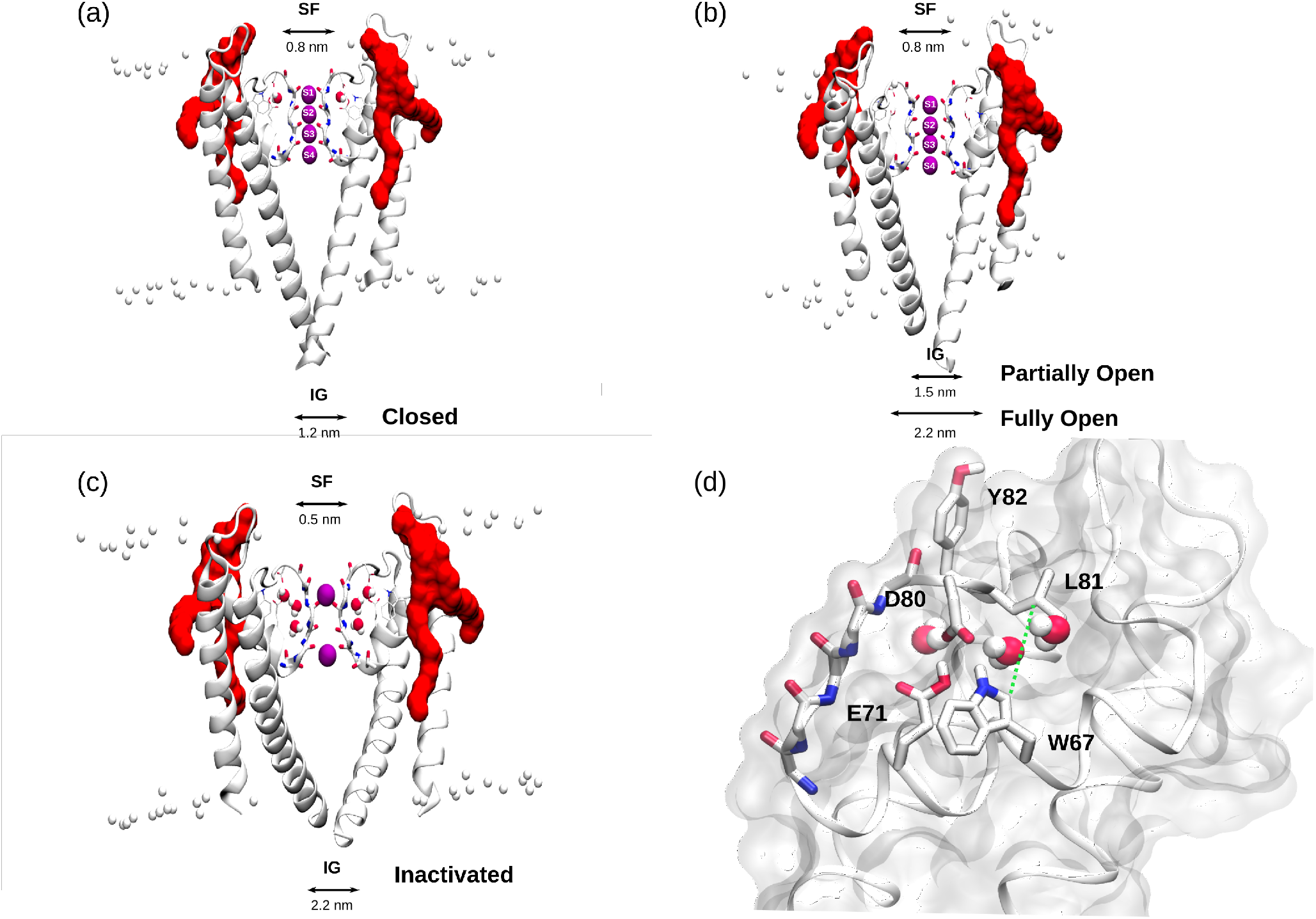
Structure of two opposing subunits of the KcsA tetramer representing the closed state (a), the partially open and fully open states (b) and, the inactivated state (c). Potassium cations are represented by purple spheres, membrane lipid headgroups by white spheres, and bound DOPG lipids by their molecular surface in red. The typical values for their inner gate (IG) and selectivity filter (SF) openings are shown below each snapshot. (d) Representative MD snapshot of the pathway taken by water molecules before the final step of inactivation. This pathway passes between L81 and W67 (green dashed line) allowing for two additional water molecules to be buried behind the selectivity filter.

The inactivation process has also been shown to be modulated by anionic lipids (***Alvis et al., 2003***; ***Demmers et al., 2003***; *Marius et al., 2008*; *Weingarth et al., 2013*; *Poveda et al., 2019*; *Zhang et al., 2021*; *Westerlund et al., 2020*). Mutagenesis experiments show that removing the co-purifying PG lipids bound between KcsA intersubunit pockets prevents inactivation, hightlighting their potential role in stabilizing the open state, or preventing entry into the inactivated one (*Poveda et al., 2019*). Several experimental techniques combined with MD simulations suggest that the inactivation mechanism is determined by the interaction between these bound PG-lipids, their anchoring residues (R89 and R64) and the inactivation-determining residues known as the “inactivation triad” (W67, E71 and D80)(*Cordero-Morales et al., 2011*; *Pless et al., 2013*; *Poveda et al., 2019*).

Molecular dynamics (MD) simulations are a powerful source of mechanistic information on the atomic level and, as such, many insights about ion channels (***Flood et al., 2019***; ***Lindahl and Sansom, 2008***; ***Carnevale et al., 2021***), including KcsA (***Poveda et al., 2019***; ***Li et al., 2018***; ***Furini and Domene, 2009***; ***Kopec et al., 2018***; ***Oakes et al., 2020***; ***Pérez-Conesa et al., 2021***; ***Westerlund et al., 2020***), have been obtained with this technique. One of the key modeling choices when using MD simulation is the choice of force field that describes the interactions between particles being simulated. For KcsA, this choice is crucial since the selectivity filter’s behaviour is remarkably force field-dependent (***Kopec et al., 2018***; ***Furini and Domene, 2020***). A recent study thoroughly compared the differences observed in the system when using either the CHARMM or AMBER force fields without concluding which best reproduced experimental findings (***Furini and Domene, 2020***). This dissimilarity suggests that the free energy surfaces that the system navigates may differ depending on the force field used to describe the system.

Estimating the free energy surface of processes as slow as KcsA inactivation, requires enhanced sampling MD simulations. Although there have been a few studies of selectivity filter constriction and ion permeation using enhanced sampling simulations (***Allen et al., 2004***; ***Fowler et al., 2013***; ***Li et al., 2018***; ***Oakes et al., 2020***; ***Furini and Domene, 2009***), a simulation study of the full inactivation path, emcompassing allosteric signal transmission between the two gates, is lacking.

Having an effective method to obtain the free energy surface of inactivation would allow studying the energetic impact of introducing perturbations in the system, such as single point mutations, or removing co-purifying PG-lipids. It is also of particular interest to understand whether the fully open state (PDB-ID 5VK6, captured using mutations to stabilize this specific conformation) is physiologically relevant, given that the partially open state (PDB-ID 3FB5) was recently shown to be the dominant state in activating conditions (***Pérez-Conesa et al., 2021***) in ssNMR experiments. Indeed, simulations based on the fully open state show spontaneous constriction of the selectivity filter(***Li et al., 2018***). Strikingly, however, constriction is observed only if the CHARMM force field is used, and no such events are observed over MD simulation timescales if the AMBER force field is used instead(***Furini and Domene, 2020***). All this begs the question: is the fully open state construct (PDB-ID 5VK6) a pre-inactivated transition state-like structure rather than a truly metastable state?

Building on previous characterization of the conformational ensembles of membrane proteins (Lev et al., 2017a; ***Fleetwood et al., 2019***, 2021; ***McComas et al., 2022***), in this work, we use the string method with swarms of trajectories (***Pan et al., 2008***) to obtain the full inactivation pathway free energy surface. Based on these simulations, we propose a potential role for residue L81 in opening a gateway for water molecules to enter behind the selectivity filter. We also contrast the free energy landscapes obtained using two different force fields, CHRAMM and AMBER. Finally, we investigate the role of co-purifying lipids by computing free energy surfaces with the bound lipids removed from their binding sites. We conclude by postulating a new inactivation mechanism, excluding the fully open state, observed using the CHARMM force field that could potentially be further supported or disproven by experimenting with an L81A construct.

## Methods

### System Preparation

KcsA coordinates were prepared using CHARMM-GUI (***Jo et al., 2017***) in the closed state. Residues 25–121 of the X-ray diffraction closed state structure (PDB-ID 5VKH (***Cuello et al., 2017***)) were used for consistency with our previous work. The selectivity filter was prepared fully loaded with potassium and one buried crystallization water behind each subunit’s selectivity filter was included. All termini were NME- and ACE-capped and residues H25, E71, E118 and E120 were protonated following NMR data in activating conditions (***Pérez-Conesa et al., 2021***). Mutations in the experimental construct (Y82A, L90C and F103A) were reverted using PYMOL (***Lill and Danielson, 2011***). The protein was embedded in a 3:1 DOPE:DOPG membrane bilayer and surrounded by a 50mM KCl and 50mM NaCl solution including ∼90 TIP3P rigid water molecules per lipid and a total of 150 lipids. The co-purifying bound PG lipids found between subunits were reconstructed as DOPG molecules following the partial coordinates of the experimental construct. Two starting coordinates were prepared one with the co-purifying lipids bound (LB) and another without these lipids (noLB). The lipid bound system was simulated using AMBER14SB+SLIPID16 and CHARMM36 (***Maier et al., 2015***; ***Jämbeck and Lyubartsev, 2012***, ***2013***; ***Huang and MacKerell, 2013***) and the lipid unbound system only with AMBER14SB producing a set of three simulation conditions: LB-AMBER, noLB-AMBER and LB-CHARMM. CHARMM36 was used rather than the more recent CHARMM36m (***Huang et al., 2016***) to enable a direct comparison with Li et al ***Li et al. (2018)***. Nevertheless, our results are consistent with published data using CHARMM36m (***Kopec et al., 2018***; ***Furini and Domene, 2020***). If unspecified, the system is assumed to have the PG lipids bound.

### Molecular Dynamics Simulations

All molecular dynamics simulations were performed using GROMACS2020.5. The NPT ensemble was simulated using the Bussi thermostat (tau=1ps T=290K) and Parrinello-Rahman barostat (𝜏 = 5 ps and P = 1 bar)(***Bussi et al., 2007***; ***Parrinello and Rahman, 1981***). The temperature was chosen for compatibility with solid state NMR conditions and our previous work. The systems were equilibrated using an extended CHARMM-GUI protocol. Additional technical details can be found in the freely available input files (https://github.com/delemottelab/KcsA_string_method_FES).

### String Method with Swarms of Trajectories

In order to sample the inactivation process, we used the string method with swarms of trajectories (***Pan et al., 2008***). Previous studies of pentameric ligand-gated ion channel activation (***Lev et al., 2017b***), GPCR activation (***Fleetwood et al., 2021***, ***2019***) and sugar porters (***McComas et al., 2022***) have established this method as a useful tool for simulating and understanding protein conformational change. This method takes as input an approximate path connecting the initial state to the final state. Then, a set of initial structures along the path are taken as beads connected by a multidimensional string representing the structural transition. The multidimensional string is parametrized by a set of generalized coordinates known as collective variables (CVs) that describe the transition path and the slow degrees of freedom of the process. From each of the beads, a set of short “swarm” unbiased simulation replicas are launched. The average drift of the swarms in the string-CVs is calculated and used to update the position of the string. Then, equilibration simulations are launched from every bead restraining the CV values of the simulation to the new string iteration thus updating the coordinates of the beads to the new string iteration. This process is repeated until the string converges and oscillates around a fixed path in CV-space. This path corresponds to a minimum free energy pathconnecting the starting and final states. Many additional iterations are run in order to intensively sample the minimum free energy path for analysis.

To obtain the initial string, a steered simulation was performed starting from the equilibrated closed state (PDB-ID 5VKH), to the partially open state X-Ray coordinates (PDB-ID 3FB5), to the fully open state (PDB-ID 5VK6) and finally to the inactivated state (PDB-ID 5VKE). A harmonic potential was applied on a set of 60 steering-CVs (Table 1) moving the target CV value at a constant speed to force the channel to go through the mentioned conformational states in a total simulation time of ∼500ns. The speed of steering was decreased in the final stages to allow the inactivation waters to enter the cavities behind the selectivity filter. The steering CVs were determined by structure comparison and bibliographical information of the system. For all steering simulations, unbiased simulations from the final coordinates were launched to check if the final state CVs remained stable and the system did not return to a fully-open-like state.

KcsA inactivation encompasses both rigid body motions such as the opening of the inner gate helical bundle and residue level motions such as the constriction of the selectivity filter, the motion of pore helix residues to stabilize it and accommodate the two additional buried water molecules per subunit. Thus to parametrize the string we used 36 interatomic distances, 11 of which are not symmetry related CVs. We chose: 18 CVs of TM2 cross- and adjacent-subunit distances to model inner gate opening, 8 CVs connecting allosteric contacts between the water cavity region and the SF (***Li et al., 2018***) that transmit the allosteric signal from the inner gate to the selectivity filter and 10 CVs related to the constriction of the SF. These CVs are a subset of the steering CVs and a result of filtering steering CVs that were unimportant in preliminary string simulation trials. The set of 36 CVs chosen to parametrize the string can be found in Table 2.

We observed in preliminary steering simulations of the CHARMM system that going from the fully open state (PDB-ID 5VK6) to the inactivated state (PDB-ID 5VKE), the contact between the sidechains of L81 and W67 spontaneously broke, in absence of any external bias explicitly acting on the distance between these two residues. This opened a gateway for two water molecules to enter the cavity behind the selectivity filter, eventually resulting in the obtention of the inactivated state with three buried waters per subunit, and an intact L81-W67 contact (***Figure 1e***). Subsequently adding this contact distance to the steering CV set promoted water penetration in all simulation conditions. In contrast, if this CV was not used during steering, water penetration was not observed until the spontaneous breaking of the mentioned contact at a significantly longer timescales, if this event even happened at all. We thus elected to include the breaking and reformation of this contact during the final inactivation step in our initial steering simulation, and included the distance between this residue pair as a string CVs.

The string method with swarms of trajectories was run using 18-bead strings obtained from the steering simulations fixing the endpoint beads. At each iteration, for every bead, a 100ps restrained simulation was performed and from each final snapshot, 32 10ps-swarms were launched to calculate the average drift of the strings. A total of ∼800 iterations were run for minimum free energy path sampling after string convergence which occurred at 100 to 300 iterations. The convergence of the string equilibrium positions was evaluated with the time evolution of the beads and the root mean squared deviation of the string with respect to the initial string (***Figure 2—figure Supplement 1*** and ***Figure 2—figure Supplement 2***). The string simulations were performed using our own string method with swarms of trajectories python package which is freely available online (https://github.com/delemottelab/string-method-swarms-trajectories).

### Free Energy Surface Calculation

Free energy surfaces were calculated using reweighted kernel density estimations of the probability distribution of CVs calculated per swarm trajectory snapshot. The weights of each swarm snapshot were obtained using a reversible maximum likelihood Markov State Model (MSM) with the deeptime python library (Hoffmann et al., 2021). The MSM was parametrized using states obtained from k-means clustering of a TICA dimensionality reduction of the string-CVs of the swarm trajectories. It is worth noting that since weights are assigned to the particular trajectory snapshot, the reweighted free energy surface can be projected to non-string CVs. Considering this, in order to have a more interpretable view of the path energetics, a one-dimensional projection of the free energy surface was also calculated. The CV chosen to describe the inactivation path is the progression along the converged path from the string simulations, 𝑠_𝑝𝑎𝑡ℎ_, being ∼0 in the closed state and ∼1 in the inactivated state. This path CV is calculated according to Braduardi’s definition (***Branduardi et al., 2007***) and using as the converged path the average of the last 25 strings generated. This CV indeed represents correctly the progression along inactivation as can be seen in the overlay of the average path-CV value with the IG vs SF projected free energy surface (***Figure 2—figure Supplement 3***). Additional details of the MSM and free energy surface calculation can be found in the following repository (https://github.com/delemottelab/KcsA_string_method_FES.).

200ns long unbiased molecular dynamics started from the different free energy basins were started. These simulations have the expected stability based on their starting values (***Figure 2—figure Supplement 5***). This is a good quality test to check the correct estimation of the general features of the free energy surface.

The statistical errors of the free energy surface by combining blocking decorrelation analysis (***Flyvbjerg and Petersen, 1989***) and bootstrapping (***Hub et al., 2010***) (***Figure 2—figure Supplement 4***). The swarm data is divided into 𝑛 blocks for 𝑛 = 2, 4, 8, 16, 32 and 150 bootstraps are done on the blocks. The set of bootstraps for a given 𝑛 is used to generate a distribution of free energy surface. The error is then calculated as the standard deviation of this free energy surface distribution. 𝑛 is chosen such that the average error over the free energy surface is maximized as a consequence of the time-decorrelation between the data of the blocks.

## Results

### Free energy surfaces reveal force field-dependent inactivation mechanisms

From the string method with swarms of trajectories, we obtained projections of the inactivation free energy surfaces onto the inner gate distance (IG, average cross-subunit distance between the CA of T112) and the selectivity filter distance (SF, average cross-subunit distance between the CA of G77 residues) (***Figure 2a-b***).

**Figure 2.**
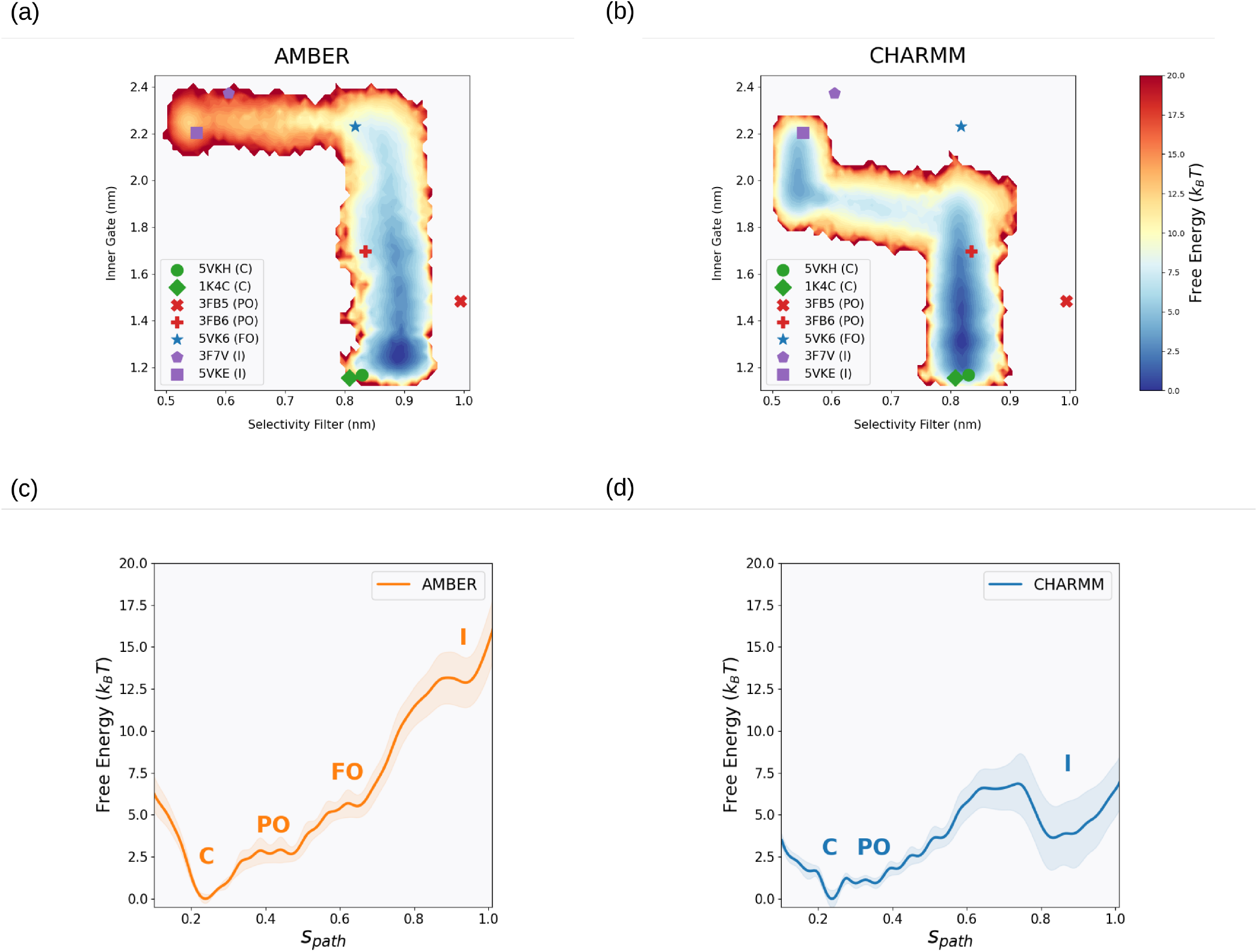
Reweighted free energy surfaces calculated from the string method with swarms of trajectories. 2D free energy surfaces projected on the inner gate distance (IG, average cross-subunit distance between the CA of T112) and the selectivity filter distance (SF, average cross-subunit distance between the CA of G77 residues) of the systems using (a) AMBER and (b) CHARMM. 1D free energy surfaces projected on the path cv (*𝑠𝑝𝑎𝑡ℎ*) of the systems using (c) AMBER and (d) CHARMM with labels denoting the state of the basin: closed (C), partially open (PO), fully open (FO) and inactivated (I). The errors associated with the 2D-free energy surface can be found in Figure 2—figure Supplement 4 **Figure 2—figure supplement 1**. Evolution of the string beads over the string method iteration (averaged over blocks of 25 iterations) and projected on the inner gate distance (IG, average cross-subunit distance between the CA of T112) and the selectivity filter distance (SF, average cross-subunit distance between the CA of G77 residues). The initial string is represented as a black dotted line. The inset plots the first ten iterations without averaging. The systems represented are (a) LB-AMBER, (b) LB-CHARMM and (c) noLB-AMBER. **Figure 2—figure supplement 2.** Evolution of the root mean squared deviation from the initial string averaged over all beads. The systems represented are LB-AMBER (orange), LB-CHARMM (blue) and noLB-AMBER (red). **Figure 2—figure supplement 3.** The weighted average value of *𝑠𝑝𝑎𝑡ℎ* projected 2D-free energy surface projected on the IG and SF. The systems represented are (a) LB-AMBER, (b) LB-CHARMM and (c) noLB-AMBER. **Figure 2—figure supplement 4.** Estimation of the uncertaintly associated with the 2D-free energy surface projected on the IG and SF (see Methods). The systems represented are (a) LB-AMBER, (b) LB-CHARMM and (c) noLB-AMBER. **Figure 2—figure supplement 5.** IG and SF projections of 200 ns MD trajectories started from representatives of different states of the inactivation cycle onto corresponding free energy surface. The starting structure is represented as a triangle. The systems represented are (a) LB-AMBER, (b) LB-CHARMM and (c) noLB-AMBER.

The free energy surface obtained using the AMBER force field (***Figure 2a***) shows an inactivation path which follows the canonical structural path inferred from X-ray crystallography data (***Doyle et al., 1998; Cuello et al., 2010a,b***, ***2017***): closed state, partially open state, fully open state and inactivated state. In addition, the IG and SF of these states take very similar values to those of the crystal structure constructs, with the exception of a ∼1 Å widening of the SF with respect to most unconstricted SF X-ray structures in the conductive states (PDB-IDs 1K4C, 5VKH, 3FB6 and 5VK6). The impact of the force field choice on the free energy surface of the systems is very strong. The CHARMM simulation explores a different minimum free energy path (***Figure 2***b). For CHARMM, the SF filter opening for the closed and partially open states is smaller than in AMBER simulations, similar to the XRD values of 0.8nm. This narrowing could rationalize the lower conductance obtained using the CHARMM force field (***Furini and Domene, 2020***). In addition, fully open-like IG distances lie outside of the minimum free energy path. In other words, using this force field, the only open state along the lowest free energy inactivation path is the partially open state. The partially open state to inactive state transition shows a correlated closing of the SF and opening of the IG, in contrast to the AMBER case where the SF constricts at a constant fully-open IG value. Additionally, the free energy barriers using CHARMM are substantially lower than using AMBER.

Our previous investigation combining solid-state NMR and MD simulations had inferred that a partially open state was more energetically favorable than a fully open state under activating conditions(***Pérez-Conesa et al., 2021***). This experimental evidence is in agreement with the lower FE basins corresponding to the partially open state obtained using both the AMBER and the CHARMM force field. Furthermore, the fact that our systems prepared with the protonation states and salt concentration used in activating conditions solid-state NMR experiments show partially open states thermodynamically accessible from the resting state exemplifies the quantitative agreement of the method with independently obtained experimental data.

The 1-D free energy surface of the path-CV, 𝑠_𝑝𝑎𝑡ℎ_, which measures the progression along the reaction path, shows profiles with comparable features using both force fields (***Figure 2***c-d). The lowest basin is the closed state (𝑠_𝑝𝑎𝑡ℎ_ ∼ 0.2). The shallow basins of the open state or open states, of higher free energy, are then explored (𝑠_𝑝𝑎𝑡ℎ_ ∼ 0.4 and 𝑠_𝑝𝑎𝑡ℎ_ ∼ 0.6 for AMBER and 𝑠_𝑝𝑎𝑡ℎ_ ∼ 0.35 CHARMM). Finally, there is a substantial free energy barrier leading to the highest free energy basin (𝑠_𝑝𝑎𝑡ℎ_ ∼ 0.9), corresponding to the inactivated state, consistent with our simulations being run under activating conditions. Energetically speaking, the main effect of the force field choice is the increase in the barrier and difference in free energy between the closed state and the inactivated state. In particular, the barrier height and difference in free energy of inactivation are ∼14kT and ∼ 13kT for AMBER and ∼6kT and ∼ 4kT for CHARMM, respectively. Given that our simulations were conducted under activating conditions, we had expected the open states to be more populated than the closed ones. Simulations carried out at higher pH may be able to resolve this inconsistency.

### Free energy landscapes offer insights into atomistic-resolution mechanistic details

Free energy surface projection along the path-CV and another CV of interest allows studying how the CV of interest evolves with the inactivation process. One can thus observe correlations between the advancement in inactivation and the CV of interest, switch-like events, or lack of correlation. A set of these projections were calculated using as CVs of interest potential allosteric communication contacts, secondary structure movements, rotamer changes, etc. (***Figure 3***, ***Figure 3—figure Supplement 1***, ***Figure 3—figure Supplement 2*** and ***Figure 3—figure Supplement 3***)

**Figure 3.**
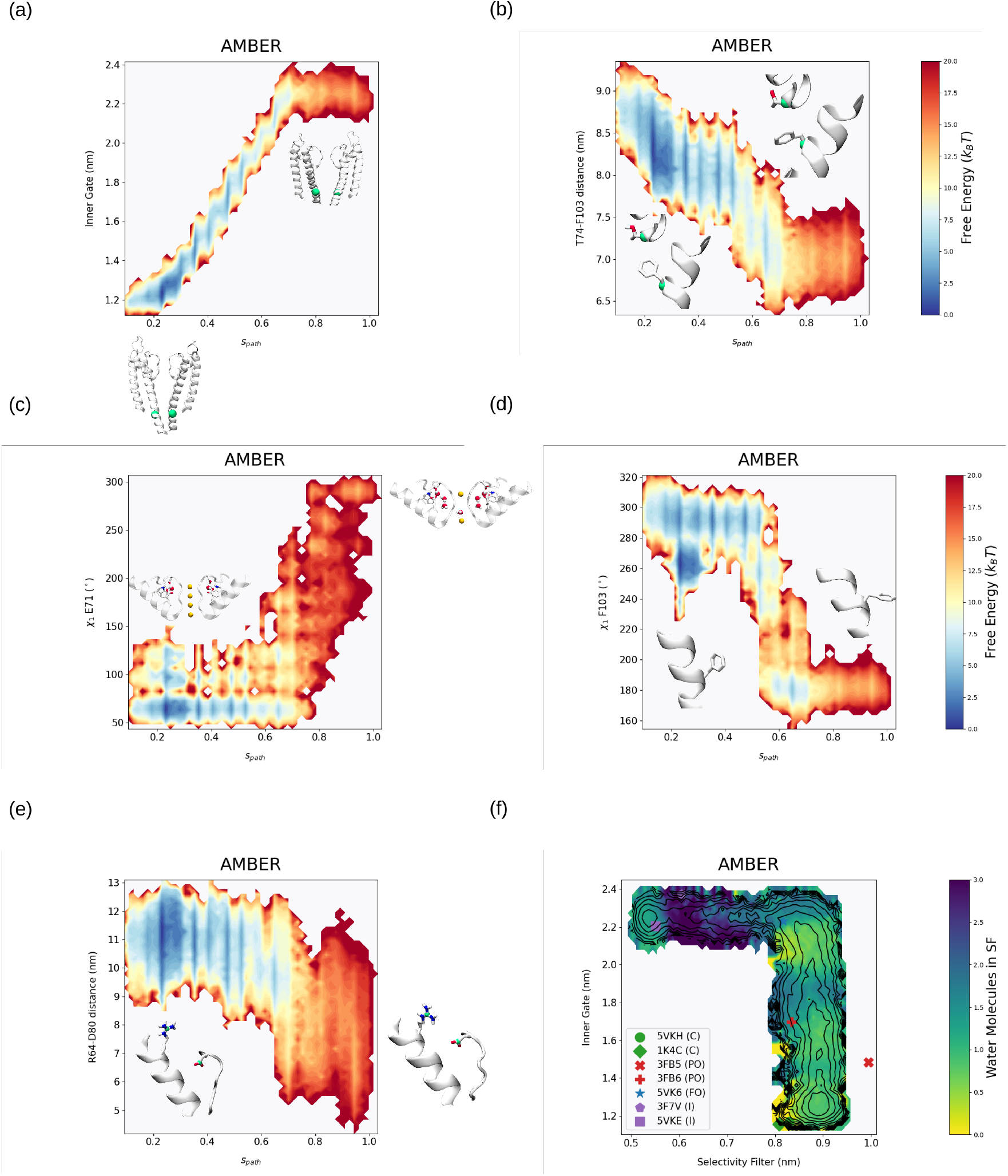
Reweighted free energy surfaces calculated from the string method with swarms of trajectories of the AMBER with lipids bound system. 2D free energy surfaces projected on the path cv (*𝑠𝑝𝑎𝑡ℎ*) and an additional CV of interest. These CVs of interest are: (a) Inner gate distance, (b) T74 CA - F103 CA distance, (c) E71 *𝜒*_1_ Janin angle, (d) F103 *𝜒*_1_ Janin angle and (e) R64 CZ - D80 CG distance. (f) Projected weighted average number of water molecules inside the selectivity filter (sites S1-S4) on the free energy surface of the AMBER lipid bound system. **Figure 3—figure supplement 1.** Reweighted free energy surfaces calculated from the string method with swarms of trajectories of the AMBER with lipids bound system. 2D free energy surfaces projected on the path cv (*𝑠𝑝𝑎𝑡ℎ*) and an additional CV of interest. These CVs of interest are: (a) T74 CA - I100 CA distance and (b) R89 CZ - D80 CG distance (c) Copurifying DOPG P - R64 CZ distance (d) Copurifying DOPG P - R89 CZ distance. **Figure 3—figure supplement 2.** Reweighted free energy surfaces calculated from the string method with swarms of trajectories of the CHARMM with lipids bound system. 2D free energy surfaces projected on the path cv (*𝑠𝑝𝑎𝑡ℎ*) and an additional CV of interest. These CVs of interest are: (a) Inner gate distance, (b) T74 CA - F103 CA distance, (c) E71 *𝜒*_1_ Janin angle, (d) F103 *𝜒*_1_ Janin angle, (e) R64 CZ - D80 CG distance, (g) T74 CA - I100 CA distance and (h) R89 CZ - D80 CG distance. (f) Projected weighted average number of water molecules inside the selectivity filter (sites S1-S4) on the free energy surface of the CHARMM lipid bound system (i) Copurifying DOPG P - R64 CZ distance (j) Copurifying DOPG P - R89 CZ distance. **Figure 3—figure supplement 3.** Reweighted free energy surfaces calculated from the string method with swarms of trajectories of the AMBER without lipids bound system. 2D free energy surfaces projected on the path cv (*𝑠𝑝𝑎𝑡ℎ*) and an additional CV of interest. These CVs of interes9toafre1:9(a) Inner gate distance, (b) T74 CA - F103 CA distance, (c) E71 *𝜒*_1_ Janin angle, (d) F103 *𝜒*_1_ Janin angle, (e) R64 CZ - D80 CG distance, (g) T74 CA - I100 CA distance and (h) R89 CZ - D80 CG distance. (f) Projected weighted average number of water molecules inside the selectivity filter (sites S1-S4) on the free energy surface of the AMBER without lipid bound system (i) Copurifying DOPG P - R64 CZ distance (j) Copurifying DOPG P - R89 CZ

In general, two types of behaviours of meaningful CVs of interest were observed: correlational and switch-like. In the first category, the minimum free energy path of the CV of interest changes linearly with the path-CV. In the second category, the FE path of the CV of interest switcheds abruptly along the path-CV. We first focus our analysis on the AMBER free energy surface, before then contrasting the differences when working with the CHARMM force field.

There is a clear correlation between path-CV and the opening of the inner gate except for the final step linking the fully open state and the inactivated one, during which it the inner gate opening remains constant (***Figure 3a***). The same behavior is observed for the contraction of the T74-F103 distance (***Figure 3b***) and the expansion of the T74-I100 distance (***Figure 3—figure Supplement 1a***), both of which are considered important contacts for the transmission of the IG signal to the SF, and (***Cuello et al., 2010a***; ***Labro et al., 2018***; ***Li et al., 2018***).

The rotameric state of E71 is a clear marker of inactivation given that for the three water molecules to be accommodated behind the SF and to form four hydrogen bonds each, it must undergo a rotameric switch (***Li et al., 2018***; ***Ostmeyer et al., 2013***). Its projected free energy surface presents a conformational distribution where E71 𝜒_1_ remains at 50-100^◦^ in closed and open states before switching to 360^◦^ during the final inactivation step (***Figure 3c***). The conformation of F103, which is known to move away from the water cavity below the SF during the inactivation process to due to the breaking of the hydrophobic contact with I100 ((***Li et al., 2018***)), displays a switch-like behavior (***Figure 3d***).

Previous experimental and computational studies have pointed out the importance of R89 and R64 interactions since they interact with the key inactivation residue D80 and also anchor the copurifying lipid, thus mediating the effect of lipid binding on inactivation (***Poveda et al., 2019***). In our simulation, we were able to point out a clear switch-like behavior for the R64-D80 distance (***Figure 3e***). The distance between the sidechains shrinks by 0.4 nm during the final step of inactivation. This highlights the involvement of R64, and indirectly of the anionic copurifying lipid in the process. We also found little to no correlation between the R89-D80 distance and the progression along the path (***Figure 3—figure Supplement 1b***) and the phosphate group of the lipids with either arginine (***Figure 3—figure Supplement 1c-d***, ***Figure 3—figure Supplement 2i-j***, ***Figure 3—figure Supplement 3i-j***,).

The 2D path-CV projections of the simulations conducted using the CHARMM force field generally follow similar patterns to those using AMBER (***Figure 3—figure Supplement 1***, ***Figure 3—figure Supplement 2*** and ***Figure 3—figure Supplement 3***). The main difference is that, for AMBER, more CVs of interest tend to adopt a switch-like behavior, while using CHARMM results in correlation-like behaviors. This follows from the nature of the last step of inactivation: using AMBER, the selectivity filter constricts at constant inner gate opening while this process is concerted in CHARMM.

To characterize the role played by the selectivity filter-bound water molecules in inactivation; we report the average number of water molecules (***Figure 3f***) in the SF along the inactivation path. The number of water molecules in the SF sites was found to be 0.5-1.5 in closed and open states. A per-site analysis of SF water occupation revealed that this corresponds mostly to water binding in S1, while sites S2 and S3 are fully loaded with K+ ions (***Figure 3—figure Supplement 8***). S4, on the other hand, transitions from being K+ bound to water bound as the channel proceeds to more and more open states. In the transition regions before the final inactivation step of SF constriction, there is an increase in average water occupation to 2-2.5 water molecules. These water molecules are localized in the S1, S2 and/or S4 sites . In the inactivated state, water mainly occupies S2. In contrast, for the CHARMM force field, the closed and conductive states are devoid of water in the selectivity filter, and S1, S2 and S4 are filled with K+ ions (***Figure 3—figure Supplement 2f***, ***Figure 3—figure Supplement 9***). In the transition region towards inactivation, the ion in S2 becomes replaced by water and S2 and S3 become hydrated in the inactivated state.

Our initial string featured the breaking and reforming of the L81-W67 contact during the final step of inactivation, allowing water molecules to bind behind the SF. This relied on the spontaneous observation of this phenomenon in the initial steering simulations (see methods). Studying the L81- W67 distance as a function of the progression of the equilibration of the string in each subunit of the channel, we can observe that, for the AMBER force field, this mechanism remains in a single one subunit(***Figure 3—figure Supplement 4***). In contrast, for the CHARMM force field, this path is followed by three of the four subunits (***Figure 3—figure Supplement 5***). This may show that this novel mechanism is energetically accessible to both force fields but only thermodynamically dominant when using the CHARMM force field. Instead, in both force fields, in the subunits where water does not enter via the L81-W67 pathway, the W67-D80 hydrogen breaks instead, leading to water penetration via this path. Surprisingly, we found no evidence of water penetration via the opening of the Y82 residue, in contrast to its purported role as a lid residue (***Ostmeyer et al., 2013***).

### Unbinding of copurifying PG-lipids stabilizes the open states

To study the effect of bound copurifying lipids on the inactivation mechanism, we prepared systems without any lipids anchored in the binding sites (noLB-AMBER simulations). During equilibration, steering and string simulations, however, randomly positioned DOPG lipids near the protein spontaneously bound or partially bound to the intersubunit binding sites. This produced a set of heterogeneous occupation states of the binding sites along the string. Nevertheless, the number of bound lipids remained lower than in the fully bound simulations, with an occupancy around one to two out of four.

Despite this, there is a clear impact of the low lipid binding site occupancy in the inactivation landscape (***Figure 4-b***): the partially open state is significantly stabilized in the absence of lipids (***Figure 4c-d***) and the fully open basin appears unstable as unbiased MD simulations initiated in this basin spontaneously convert to partially open states (***Figure 2—figure Supplement 5***). This suggests that stabilization of the partially open state by the removal of bound lipids can explain the increase in open probability and reduced inactivation of the channel upon mutation of the PG-anchoring residues (***Poveda et al., 2019***).

**Figure 4.**
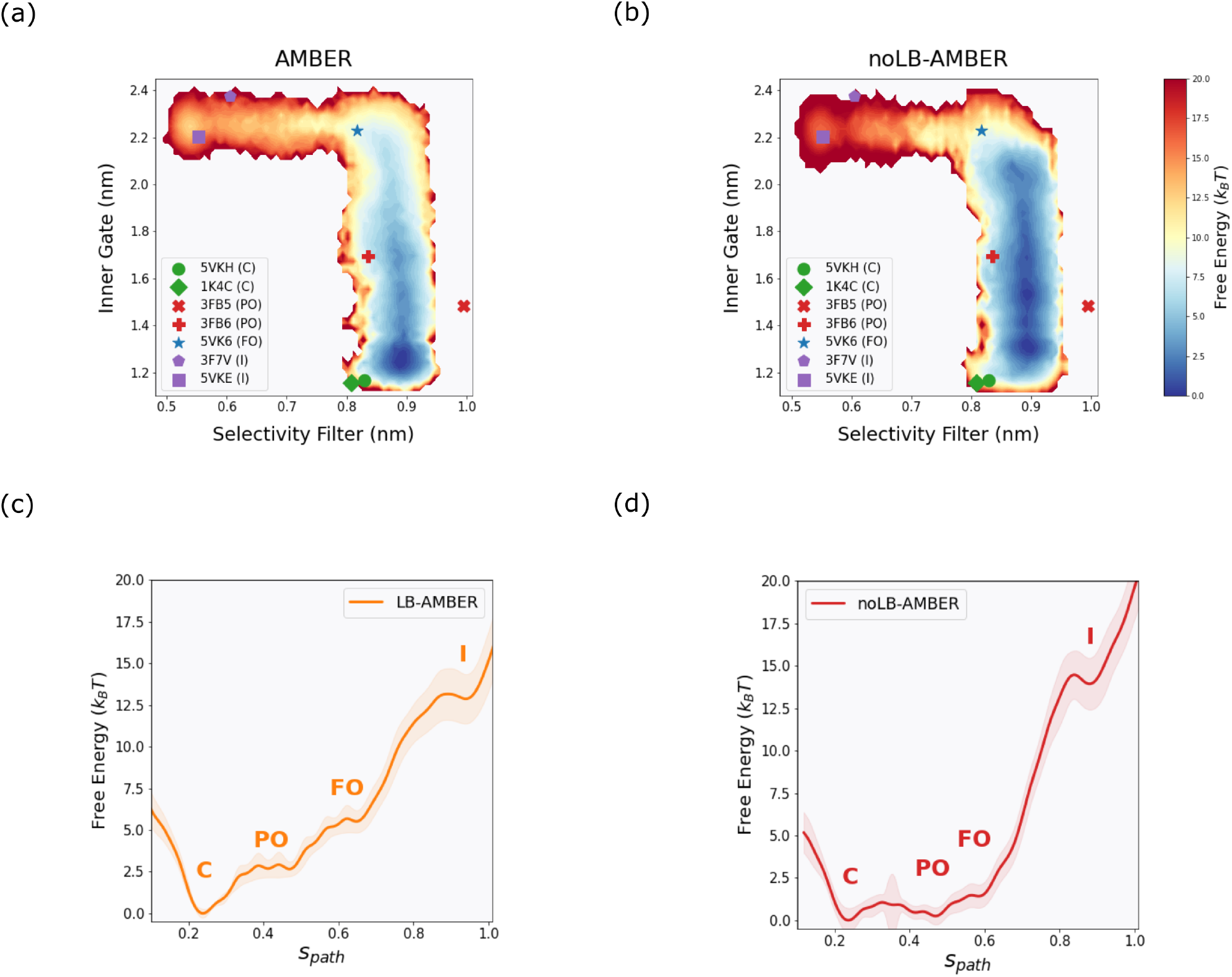
Reweighted free energy surfaces calculated from the string method with swarms of trajectories. 2D free energy surfaces projected on the inner gate distance (IG, average cross-subunit distance between the CA of T112) and the selectivity filter distance (SF, average cross-subunit distance between the CA of G77 residues) of the system with (a) PG-lipids bound and (b) no PG-lipids bound using the AMBER force field. 1D free energy surfaces projected on the path cv (*𝑠𝑝𝑎𝑡ℎ*) of the system with (a) PG-lipids bound and (b) no PG-lipids bound using the AMBER force field with labels denoting the state of the basin: closed (C), partially open (PO), fully open (FO) and inactivated (I).

## Discussion and conclusion

Molecular dynamics simulations offer atomistic insights into the function of biomolecules and can complement other experimental techniques advantageously. One of their main drawbacks is that the timescales that they allow to sample are generally much shorter than phenomena of physiological or even biophysical relevance. Enhanced sampling MD simulations have emerged as a way to expand the scope of MD simulations, while keeping an atomistic level of resolution. Many such techniques require a reduction of the dimensionality of the problem via an identification of a low number of degrees of freedom, or collective variables, that can separate different functional states of the system. Such dimensionality reduction is non trivial, and may not even be possible for all systems and processes. We have thus chosen a variant of the string method, called the string method with swarms of trajectories, in which a single minimum free energy paths linking conformational states is optimized. This method doesn’t require a dimensionality reduction as drastic as other methods, since the minimum free energy path can be optimized in high-dimension. The resulting conformational ensemble can later be projected along any collective variable interest, resulting in low-dimension free energy landscapes that can be interpreted.

Here, we have calculated the inactivation free energy surface of KcsA. These landscapes, calculated under activatory (low pH and high salt concentration) conditions, show stable closed and partially open states. The fully open state appears less stable. The pathway to inactivation features a relatively high free energy barrier. Enhanced sampling enables us to characterize the entire inactivation path under activating conditions, including the inactivated state, even if this state is not energetically favorable and would only be seldom visited using regular MD simulations. Conformational free energy landscapes are typically obtained aggregating several hundreds of microseconds of MD simulations data. Using this methodology represents a substantial computational gain, such that this type of study can be extended to study the effect of mutations, changes in pH or ion concentration.

Another major issue one is faced when using MD simulations is the choice of a potential interaction energy function (or force field) that models the processes of interest with enough accuracy. In this case, contrasting the landscapes obtained using two popular force fields suggests diverging mechanisms for the most energetically favorable inactivation cycle of KcsA. Using AMBER suggests that the fully open state is the last metastable state before selectivity filter constriction (***Figure 5a***). In this case, the selectivity filter constraints at constant inner gate opening. In contrast, using CHARMM reveals that the partially open state is the final metastable state before selectivity filter constriction, the fully open state lying outside the minimum free energy path (***Figure 5b***). In this case, the selectivity filter constricts concertedly with an inner gate expansion. This mechanism defies the current canonical inactivation path of KcsA. Interestingly, this mechanism (***Figure 5b***) is obtained despite the initial string preparation assuming the canonical path (***Figure 5a***).

**Figure 5.**
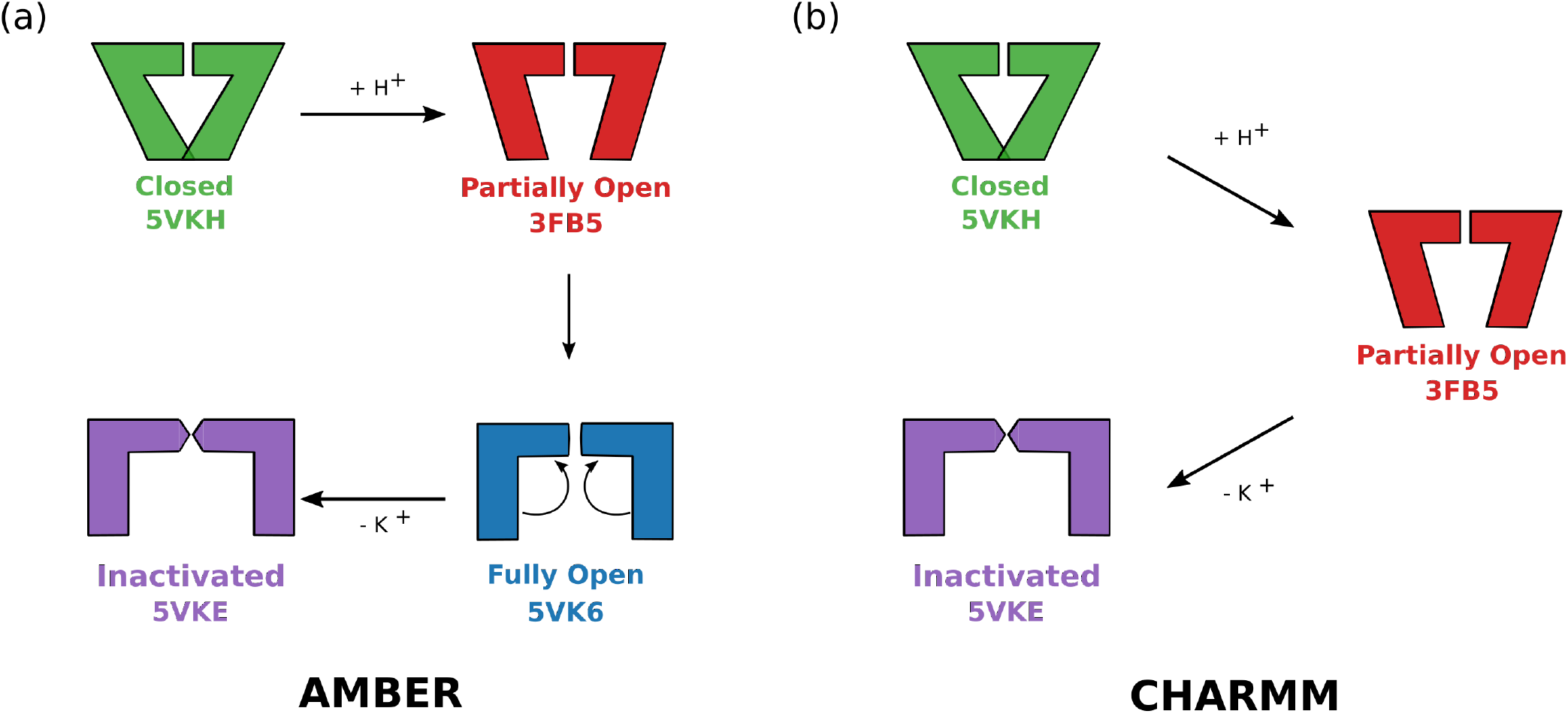
Schematic depiction of the two possible inactivation mechanisms of KcsA as described by the minimum free energy paths obtained by the string method with swarms of trajectories for the AMBER (a) and CHARMM (b) force fields.

Molecular dynamics simulations started from the fully open state derived from the XRD structure with PDB-ID 5VK6 have been repeatedly observed to leads to selectivity filter constriction when the CHARMM force field is used (***Ostmeyer et al., 2013***; ***Li et al., 2018***; ***Furini and Domene, 2020***). This phenomenon is even observed for the constitutively conductive mutant E71A (***Furini and Domene, 2020***). In contrast, the AMBER force field does not lead to inactivation of the WT channel on the MD timescale (***Furini and Domene, 2020***). It has been argued that the extremely fast kinetics of this process in CHARMM simulations contrasts with its slower nature in experiments (seconds timescale) because this process represent fast SF switching instead of inactivation (***Jekhmane et al., 2019***). Others have argued that this is rather due to a bias of the force field towards the inactivated state (***Furini and Domene, 2020***). Our free energy surfaces paint a slightly more complex picture. For the CHARMM force field, states resembling 5VK6 would lie in shallow high-free basins outside the minimum free energy path. In this way, the force field does not bias towards inactivation directly but rather displays a different inactivation path excluding 5VK6-like structures. The less favorable pathway passing through the fully activated state presumably features a spontaneous downhill process linking the fully open and the inactive state. The AMBER force field, on the other hand, does incorporate a fully-open state in its mean free energy path. These two force fields differ in several ways, and precisely pinpointing the reason(s) for which these different pathways are observed would require a separate study. Likely candidates leading to different mechanisms are different ion-carbonyl and ion-water interaction strengths (***Kopec et al., 2018***; ***Furini and Domene, 2020***), as well as more recently highlighted differences in the parametrization of glutamate charge and dihedral side chains (***Kopec et al., 2022***). Nevertheless, both force fields yield inactivation barriers that are orders of magnitude lower than what is expected from electrophysiology experiments (***Ostmeyer et al., 2013***) if we assume a transition state theory approximation of the kinetics (Equa-tion 1).

Different experimental evidence offer more support to one or the other mechanism we observe. The AMBER free energy surface follows the X-diffraction structures and displays a higher inactivation barrier. The CHARMM force field, on the other hand, results in landscapes in agreement with fact that the one of the dominant states in activating conditions is the partially open state, as revealed by a combination of ssNMR+MD.

The block of conduction during inactivation appears to result from pinching at the selectivity filter, which is triggered by water molecules entering the selectivity filter, and the cavities behind the TVGYG sequence. These phenomena appear to follow opening of the inner gate, which seems to allosterically trigger changes in the region surrounding the SF. The contact between the residue at the bottom of the selectivity filter, T74, and the hydrophobic residues on the pore lining residues I100 and F103 have previously been suggested to be important (***Cuello et al., 2010a***; ***Labro et al., 2018***; ***Li et al., 2018***). Interestingly, our landscape show that T74 and I100 gradually move away from one another along the opening pathway while T74 and F103 gradually move closer together. Interestingly, the sidechain of F103 keeps its upward orientation during the first opening steps of the inner gate, and abruptly undergoes a change to a horizontal conformation after passing through the partially open state, likely enabling a tighter contact between T74 and F103. The two force fields we used in this work are broadly consistent in this description. However, the last step towards the inactivated state happens at constant inner gate opening in AMBER while it happens as the gate further opens in CHARMM. All the degrees of freedom we have probed reveal a similar behavior with abrupt transitions in AMBER and more gradual ones in CHARMM.

Our simulations are consistent with selectivity filter pinching been triggered by the penetration of water molecules in the selectivity filter, as previously observed by different research groups (***Furini and Domene, 2020***; ***Kopec et al., 2019***). While both force fields paint a different detailed mechanistic picture, they are broadly consistent in reporting that replacement of a K+ ion by a water molecule in S2 is a key step towards destabilization of the SF in its conductive conformation, in agreement with recent observations from ssNMR (***Rohaim et al., 2022***).

Y82 was first proposed to be the “lid” that had to open to allow the buried water molecules to enter the cavity behind the SF on the basis of unbiased MD simulations of the closed state using the CHARMM force field (***Ostmeyer et al., 2013***). This same observation in fully-open-like state simulations (***Li et al., 2018***) and the fact that the Y82A mutant displays fast inactivation solidified this hypothesis. Our simulations show that for the CHARMM force field, the fully-open like states lie outside the minimum free energy path potentially, making simulations initiated in this state of limited relevance. In our simulations, the sidechain 𝜒_1_ and 𝜒_2_ of Y82 show little correlation with the path-CV in nearly all subunits analyzed. Here, we instead find that water entry proceeds via a pathway that is opened thanks to the breaking of the L81-W67 or the D80-W67 contacts. An alternative explanation for the fast inactivation of the Y82A mutant could be that removing the bulky tyrosine sidechain facilitates the opening of a water pathway adjacent to this residue, possibly involving W67. Mutating L81 could potentially add clarifying evidence to this question. Indeed, our research suggests that the mutation would enhance inactivation by allowing water to enter behind the SF without needing to wait for the L81-W67 gateway to open. In addition, given the strong force field dependence of this phenomenon, this information could help in determining which force field better describes the system in general: since L81 pathway is correlated with the overall inactivation mechanism, investigating L81A could shed light on the question whether if the fully open state is a part of KcsA’s inactivation cycle. Indeed, if the L81A mutation conserves wildtype inactivation, L81 would most likely not be involved in the mechanism suggesting that the AMBER mechanism is most likely. On the other hand, if L81A is a fast inactivator, this evidence would support the CHARMM mechanism which excludes the fully open-like states and highlights the importance of breaking the L81-W67 contact for water penetration.

To gain insights into the effect of bound POPG lipids on the inactivation mechanism, we have compared our original free energy surfaces, in which co-purifying lipids were indeed bound to the channel, with systems in which these lipids were removed. We observed that the removal of these lipid molecules stabilizes partially open states, potentially offering an explanation for the increased open probability observed in single-channel recordings of R64A and R89A mutants. These mutations are indeed proposed to lead to a destabilization of lipid binding. We could also observe that D80 and R64 come closer together as inactivation proceeds. When no lipids are bound, this stabilizing interaction appears less likely to form, which probably leads to a less likely destabilization of the interactions that keep the SF in a non-pinched, conductive, state.

KcsA has long been thought of as a prototype for K+ channel function. Recent evidence is however pointing towards the fact that its inactivation mechanism may be quite unique. Indeed, new structures of the fast inactivating W434F mutant show that the Drosophila Shaker channel likely inactivates via a different C-type inactivation mechanism, which involves widening of the upper part of the SF instead of pinching at the first Glycine residue (***Tan et al., 2022a***). This appears to be triggered by the destabilization of the network of interactions behind the selectivity filter (***Tan et al., 2022b***), triggered by the breaking of the D447-W434 (KcsA numbering D80-W67) contact following the mutation of W434 to Phe (***Pless et al., 2013***). This leads to a repositioning of the entire loop of residues that follow the TVGYG sequence, with a notable reorientation of D447 in a solution-facing position. A similar mechanism was described in structures of Kv1.3 (***Selvakumar et al., 2022***), of the Kv1.2 W362F mutant (***Reddi et al., 2022***), and could also be at play in K2P channels (***Lolicato et al., 2020***). In KcsA, on the other hand, this work agrees with previous suggestions that the breaking of D80-W67, when it occurs, does not lead to a major reorientation of the loop containing D80, but instead to the pinching of the SF stabilized by water molecules that were able to enter cavities via the pathway made accessible by the breakign of this contact.

A singularity in KcsA is the presence of an aspartic acid in position E71. This residue, found to be protonated by ssNMR, interacts with D80 and W67 (***Figure 1d***). We have found that E71 must adopt a different rotameric state to accommodate the water molecules bound to the back of the selectivity filter in the inactivated state. However, in our simulations, the orientation of the E71 side chain remains the same throughout the conformational change, namely E71 stays in an upward facing position even in the inactivated state. Other K+ channels generally feature a hydrophobic residue at this position. MthK, for example, contains a Val residue, V55. The presence of an aspartate residue in this position is thought to favor the stabilization of water molecules behing the selectivity filter. MD simulations of the V55E, engineered to mimic KcsA, reveal that this mutant appears to inactivate through a mechanism more similar to Shaker than to KcsA(***Kopec et al., 2022***), though SF carbonyl flipping is reminiscent of the behavior in KcsA(***Boiteux et al., 2020***; ***Kopec et al., 2022***). This is accompanied by a reorientation of E71 in a horizontal orientation, a phenomenon that is never observed in our study. We note, however, that MthK features a Tyr in place of W67, Y51, and the details of the interactions between residues is thus overall different in these channels, making transposing the mechanistic insights from this study to other channels non trivial.

## Acknowledgements

S.P.-C. was supported by a postdoctoral scholarship of the Gustafsson foundation. L.D. would like to thank the support of Science for Life Laboratory, the Göran Gustafsson Foundation, and the Swedish Research Council (Grants No. VR-2018-04905, 2019-02433 and 2022-04305). Simulations were performed on computational resources provided by the Swedish National Infrastructure for Computing (SNIC) at the PDC Centre for High Performance Computing (PDC-HPC). Additionally, the authors would like to thank Wojciech Kopec for the interesting discussions.

## Data Availability

The data and code to reproduce this work can be found at https://osf.io/snwbc/ and https://github.com/delemottelab/KcsA_string_method_FES. The python package to run the string method with swarms of trajectories is freely available at https://github.com/delemottelab/string-method-swarms-trajectories

## Transition state theory for rate estimation

In the context of transition state theory the rate of a process, 𝑘 is given by the Eyring equation:

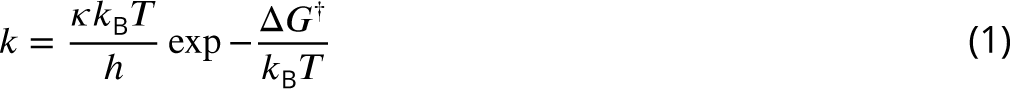

Where 𝜅 is a correction factor that is equal to 1 if transition state theory is followed ideally, 𝑘_B_ is the Boltzmann constant, ℎ is Planck’s constant, 𝑇 is the temperature and Δ𝐺^†^ is the reaction barrier. The inverse of 𝑘 is related to the mean-life of the process.

## Biased collective variables

**Appendix 0—table 1.** Distance CVs used for the steering simulations. AB and CD subunits are opposing each other and AC, CB, BD and DA subunits are side by side in a clockwise fashion observing the channel from the intracellular side.

| Residue | Atom | Residue | Atom | Sub-units |
| --- | --- | --- | --- | --- |
| 77 | CA | 77 | CA | AB, CD |
| 77 | CA | 77 | CA | AC, DB, AD, DA |
| 104 | CA | 77 | CA | AB, CD |
| 108 | CA | 77 | CA | AB, CD |
| 112 | CA | 77 | CA | AB, CD |
| 103 | CZ | 74 | CG2 | AA, BB, CC, DD |
| 100 | CD | 75 | CG2 | AA, BB, CC, DD |
| 103 | CZ | 100 | CD | AD, BD, BC, AC |
| 114 | CA | 32 | CA | AC, CB, BD, DA |
| 32 | CA | 114 | CA | CA, BC, DB, AD |
| 118 | CA | 28 | CA | AD, BD, BC, AC |
| 107 | OG1 | 101 | OG1 | AD, BD, BC, AC |
| 67 | NE1 | 80 | CG | AA, BB, CC, DD |
| 71 | OE1 | 68 | CA | AA, BB, CC, DD |
| 71 | OE1 | 78 | HN | AA, BB, CC, DD |
| 74 | CA | 79 | CA | AA, BB, CC, DD |
| 67 | CD1 | 81 | CG | AA, BB, CC, DD |

**Appendix 0—table 2.** Distance CVs used for the string simulations. AB and CD subunits are opposing each other and AC, CB, BD and DA subunits are side by side in a clockwise fashion observing the channel from the intracellular side.

| Residue | Atom | Residue | Atom | Sub-units |
| --- | --- | --- | --- | --- |
| 77 | CA | 77 | CA | AB, CD |
| 77 | CA | 77 | CA | AC, DB, AD, DA |
| 104 | CA | 77 | CA | AB, CD |
| 108 | CA | 77 | CA | AB, CD |
| 112 | CA | 77 | CA | AB, CD |
| 103 | CZ | 100 | CD | AD, BD, BC, AC |
| 114 | CA | 32 | CA | AC, CB, BD, DA |
| 32 | CA | 114 | CA | CA, BC, DB, AD |
| 118 | CA | 28 | CA | AD, BD, BC, AC |
| 107 | OG1 | 101 | OG1 | AD, BD, BC, AC |
| 67 | CD1 | 81 | CG | AA, BB, CC, DD |

**Figure 2—figure supplement 1.**
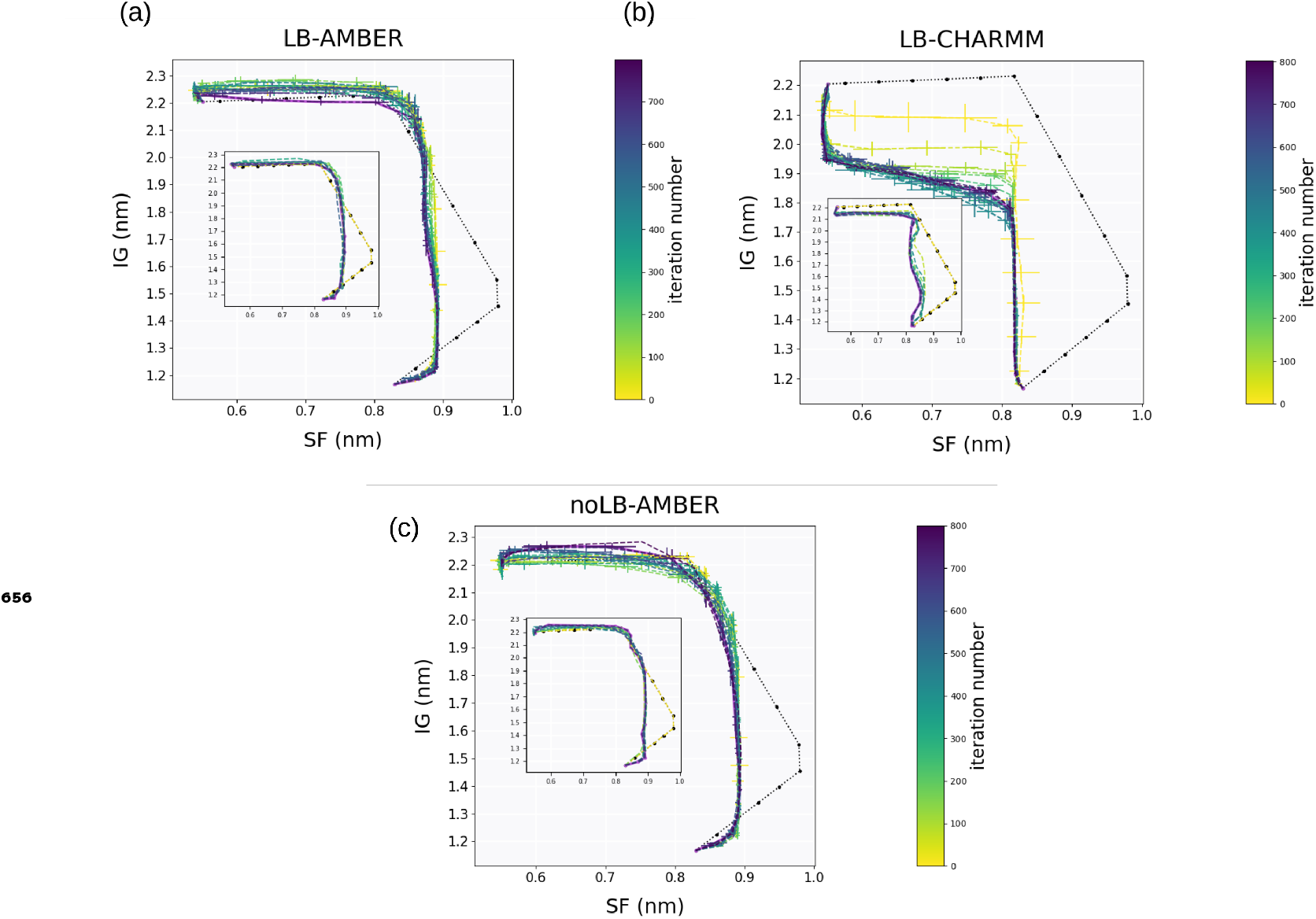
Evolution of the string beads over the string method iteration (averaged over blocks of 25 iterations) and projected on the inner gate distance (IG, average cross-subunit distance between the CA of T112) and the selectivity filter distance (SF, average cross-subunit distance between the CA of G77 residues). The initial string is represented as a black dotted line. The inset plots the first ten iterations without averaging. The systems represented are (a) LB-AMBER, (b) LB-CHARMM and (c) noLB-AMBER.

**Figure 2—figure supplement 2.**
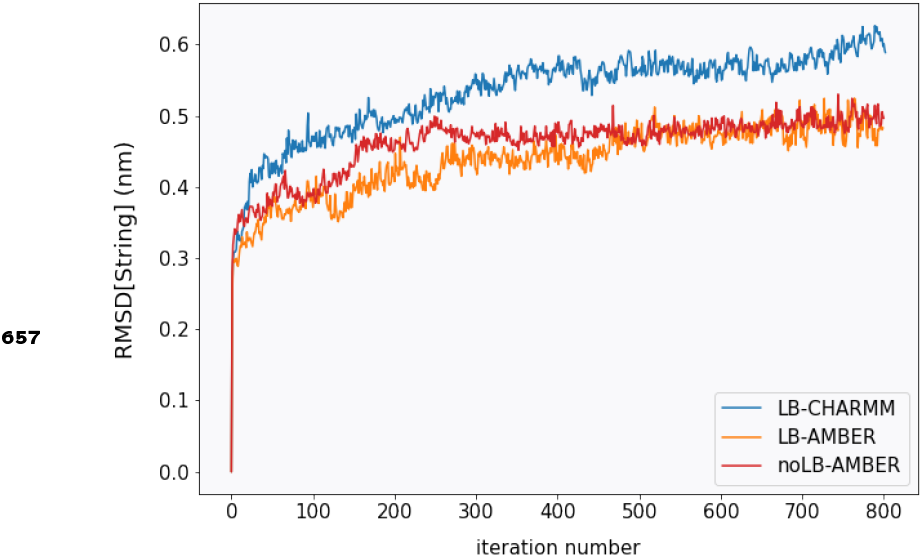
Evolution of the root mean squared deviation from the initial string averaged over all beads. The systems represented are LB-AMBER (orange), LB-CHARMM (blue) and noLB-AMBER (red).

**Figure 2—figure supplement 3.**
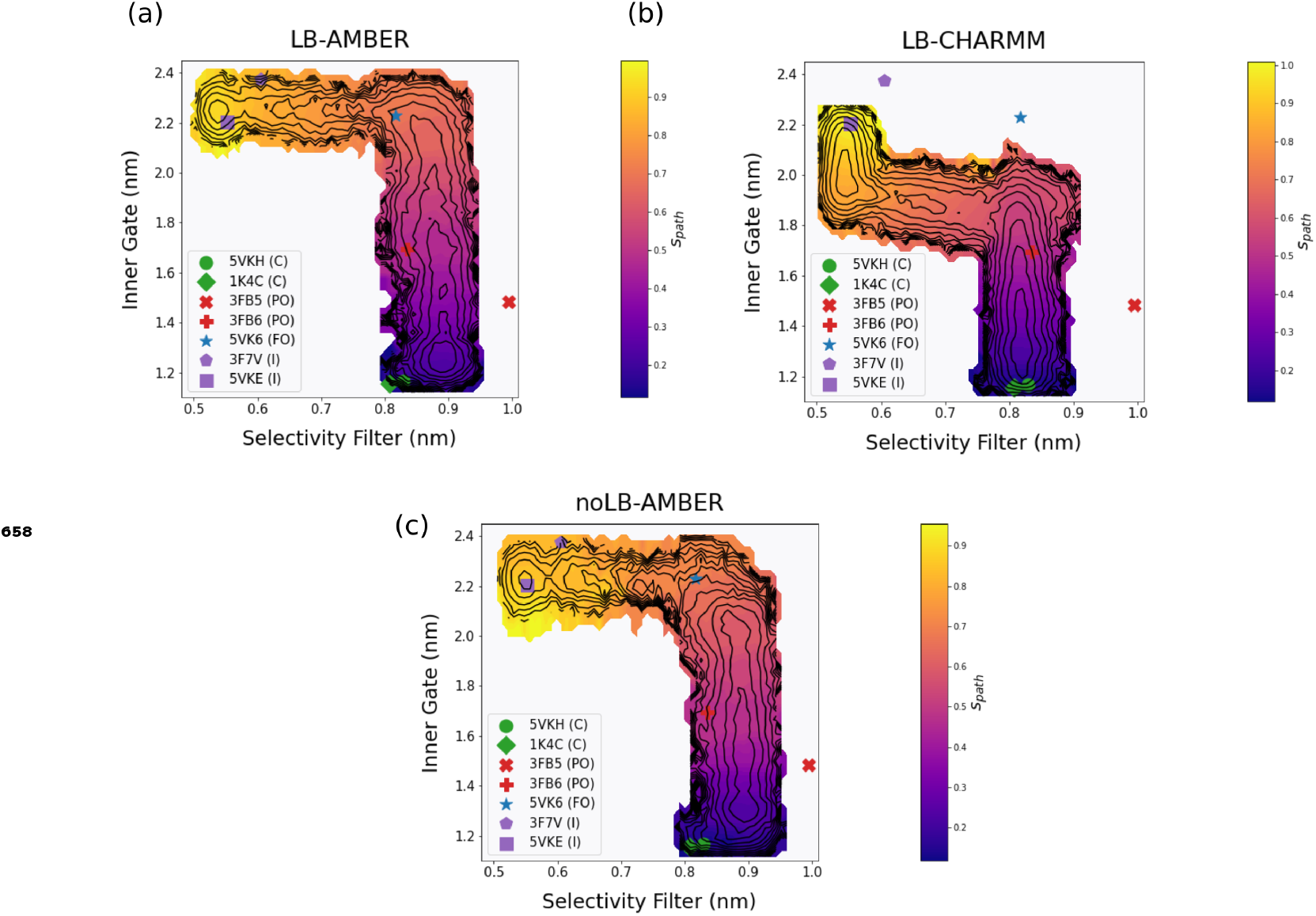
The weighted average value of 𝑠_𝑝𝑎𝑡ℎ_ projected 2D-free energy surface projected on the IG and SF. The systems represented are (a) LB-AMBER, (b) LB-CHARMM and (c) noLB-AMBER.

**Figure 2—figure supplement 4.**
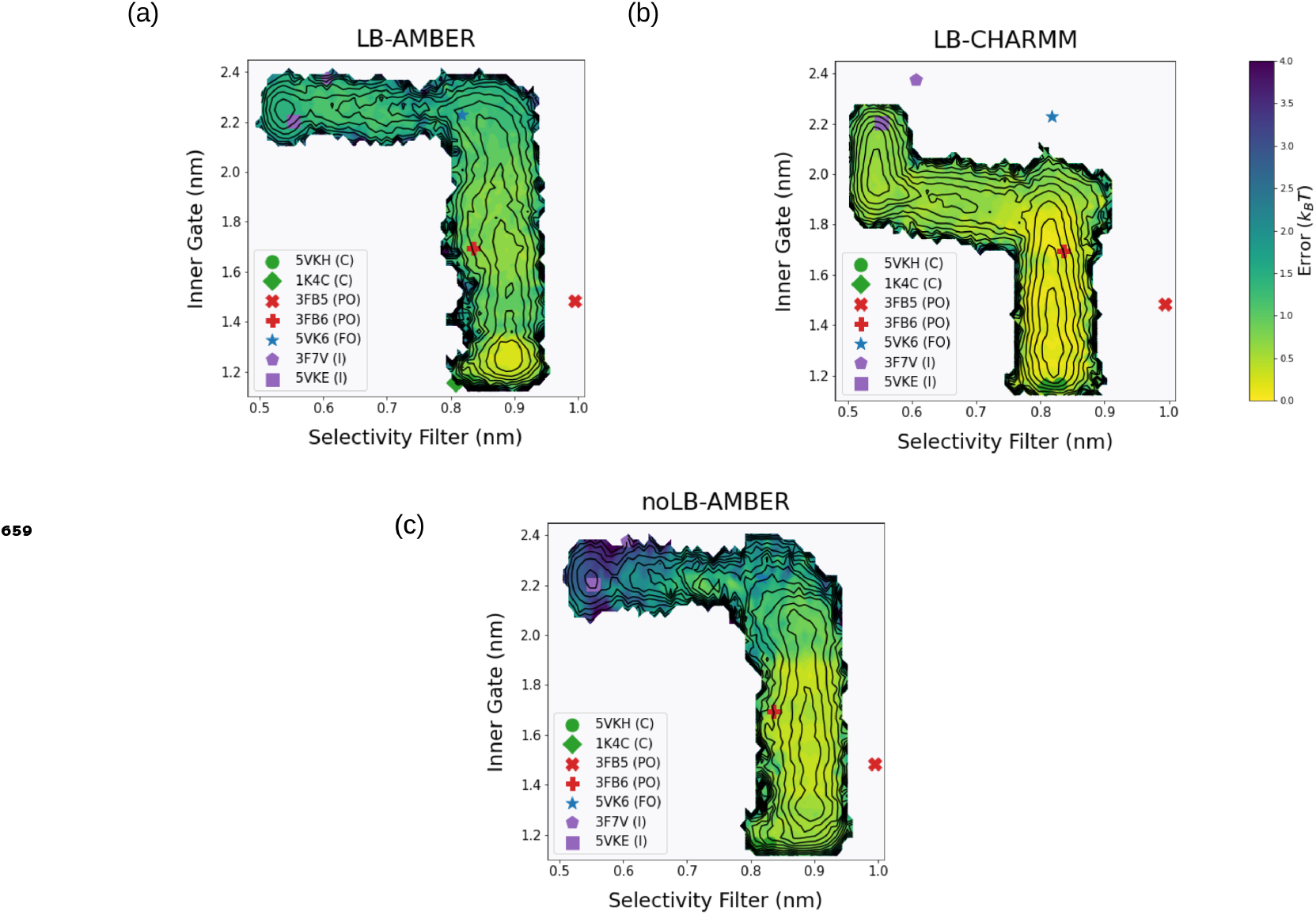
Estimation of the uncertaintly associated with the 2D-free energy surface projected on the IG and SF (see Methods). The systems represented are (a) LB-AMBER, (b) LB-CHARMM and (c) noLB-AMBER.

**Figure 2—figure supplement 5.**
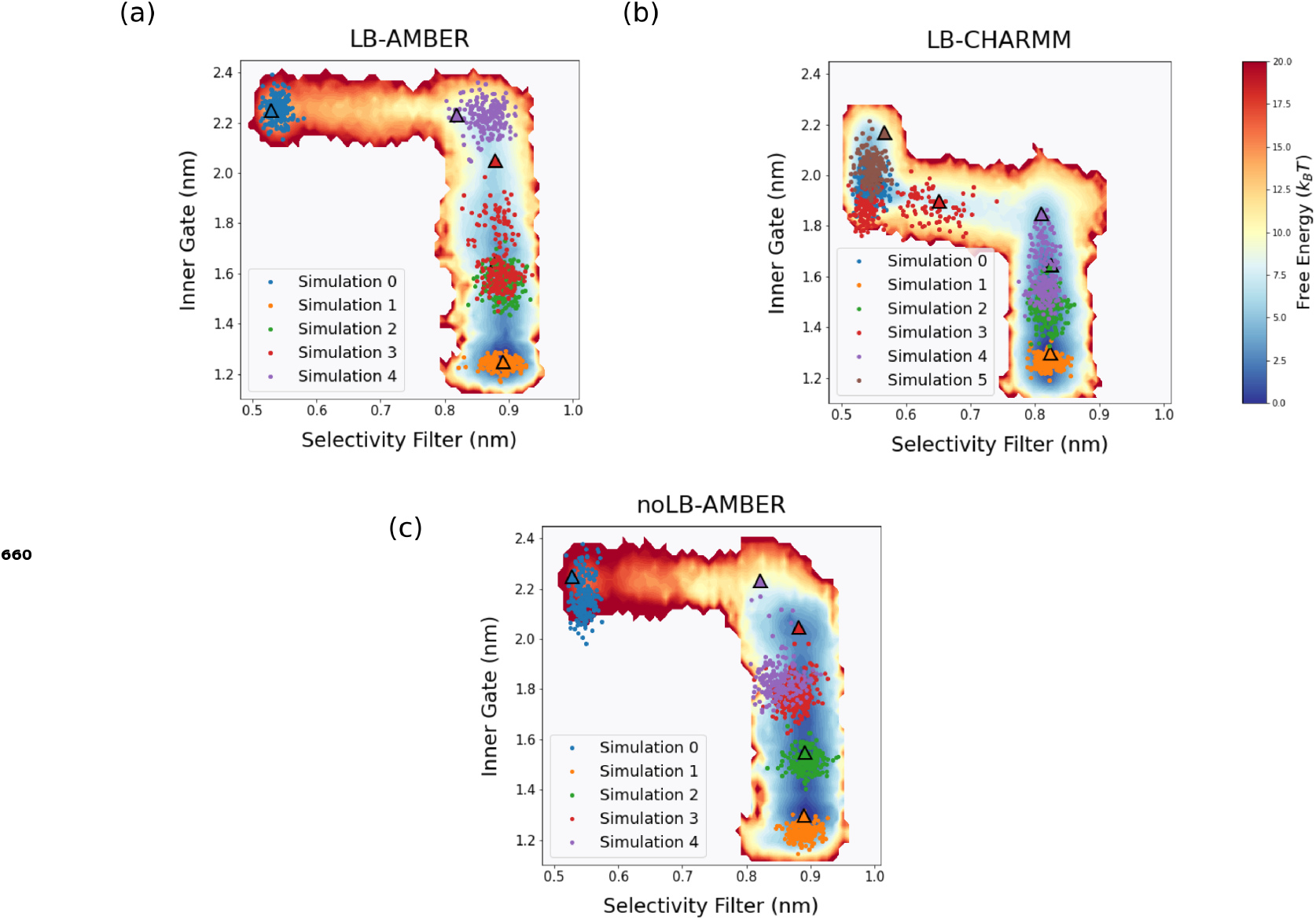
IG and SF projections of 200 ns MD trajectories started from representatives of different states of the inactivation cycle onto corresponding free energy surface. The starting structure is represented as a triangle. The systems represented are (a) LB-AMBER, (b) LB-CHARMM and (c) noLB-AMBER.

**Figure 3—figure supplement 1.**
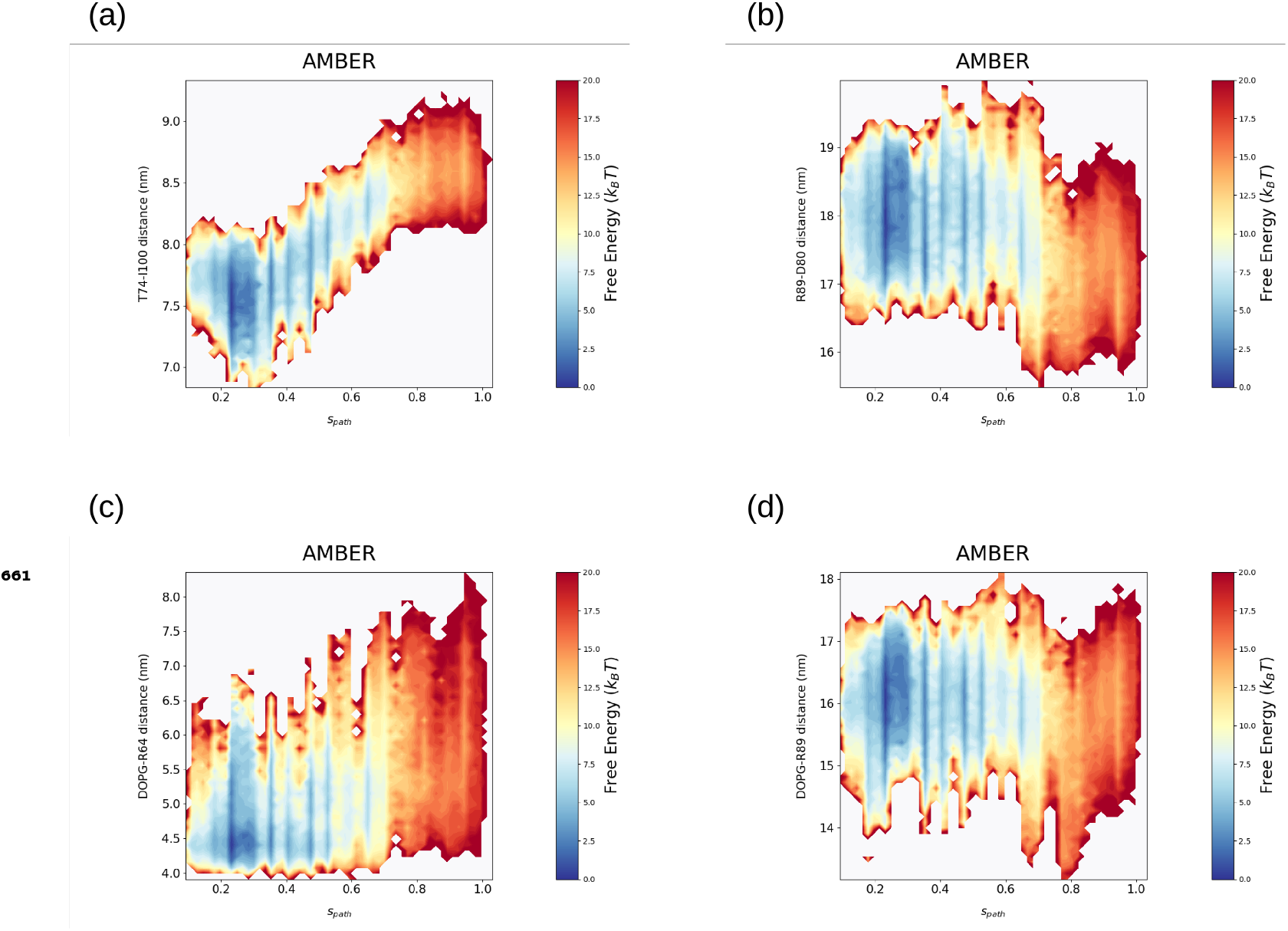
Reweighted free energy surfaces calculated from the string method with swarms of trajectories of the AMBER with lipids bound system. 2D free energy surfaces projected on the path cv (𝑠_𝑝𝑎𝑡ℎ_) and an additional CV of interest. These CVs of interest are: (a) T74 CA - I100 CA distance and (b) R89 CZ - D80 CG distance (c) Copurifying DOPG P - R64 CZ distance (d) Copurifying DOPG P - R89 CZ distance.

**Figure 3—figure supplement 2.**
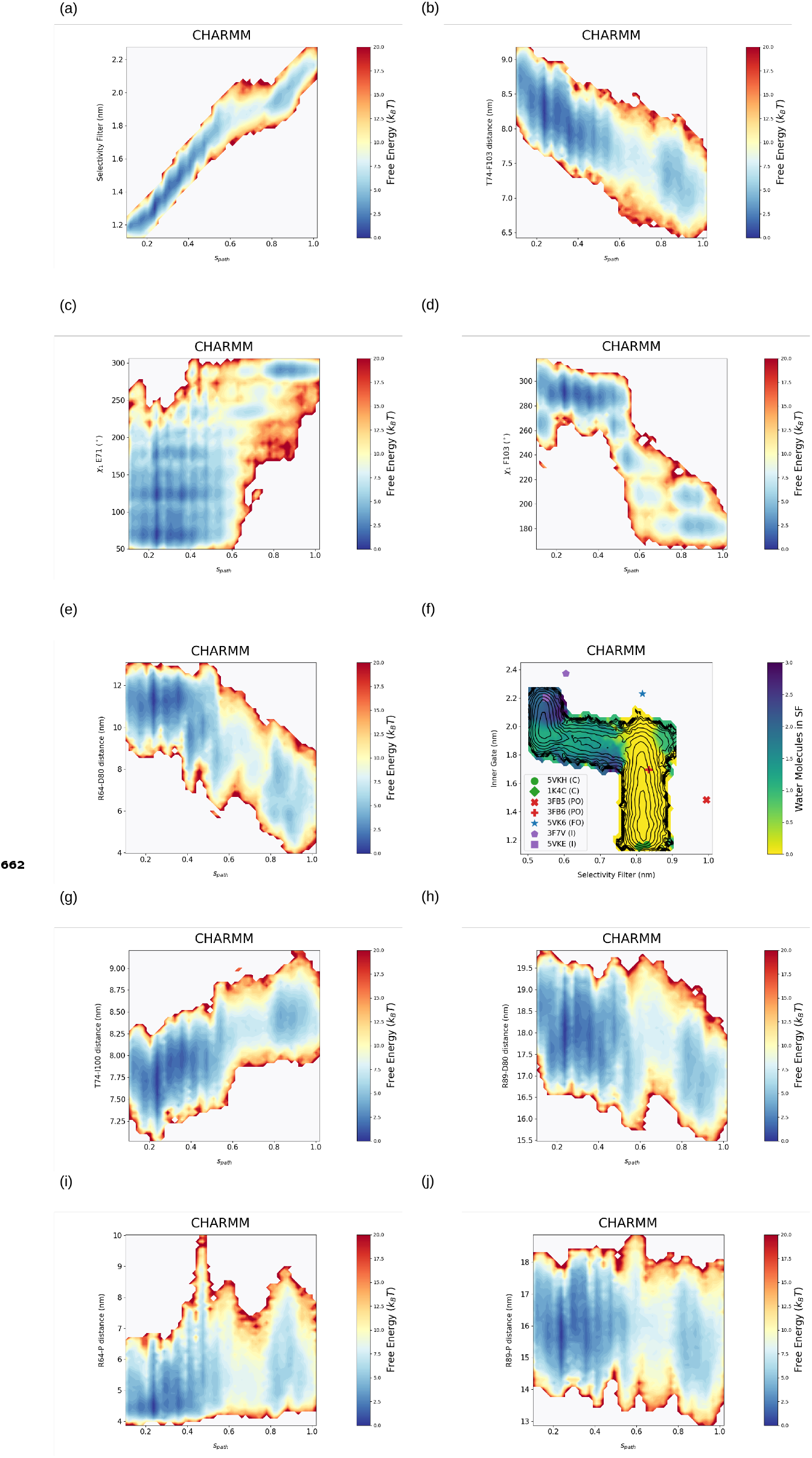
Reweighted free energy surfaces calculated from the string method with swarms of trajectories of the CHARMM with lipids bound system. 2D free energy surfaces projected on the path cv (𝑠_𝑝𝑎𝑡ℎ_) and an additional CV of interest. These CVs of interest are: (a) Inner gate distance, (b) T74 CA - F103 CA distance, (c) E71 𝜒_1_ Janin angle, (d) F103 𝜒_1_ Janin angle, (e) R64 CZ - D80 CG distance, (g) T74 CA - I100 CA distance and (h) R89 CZ - D80 CG distance. (f) Projected weighted average number of water molecules inside the selectivity filter (sites S1-S4) on the free energy surface of the CHARMM lipid bound system (i) Copurifying DOPG P - R64 CZ distance (j) Copurifying DOPG P - R89 CZ distance.

**Figure 3—figure supplement 3.**
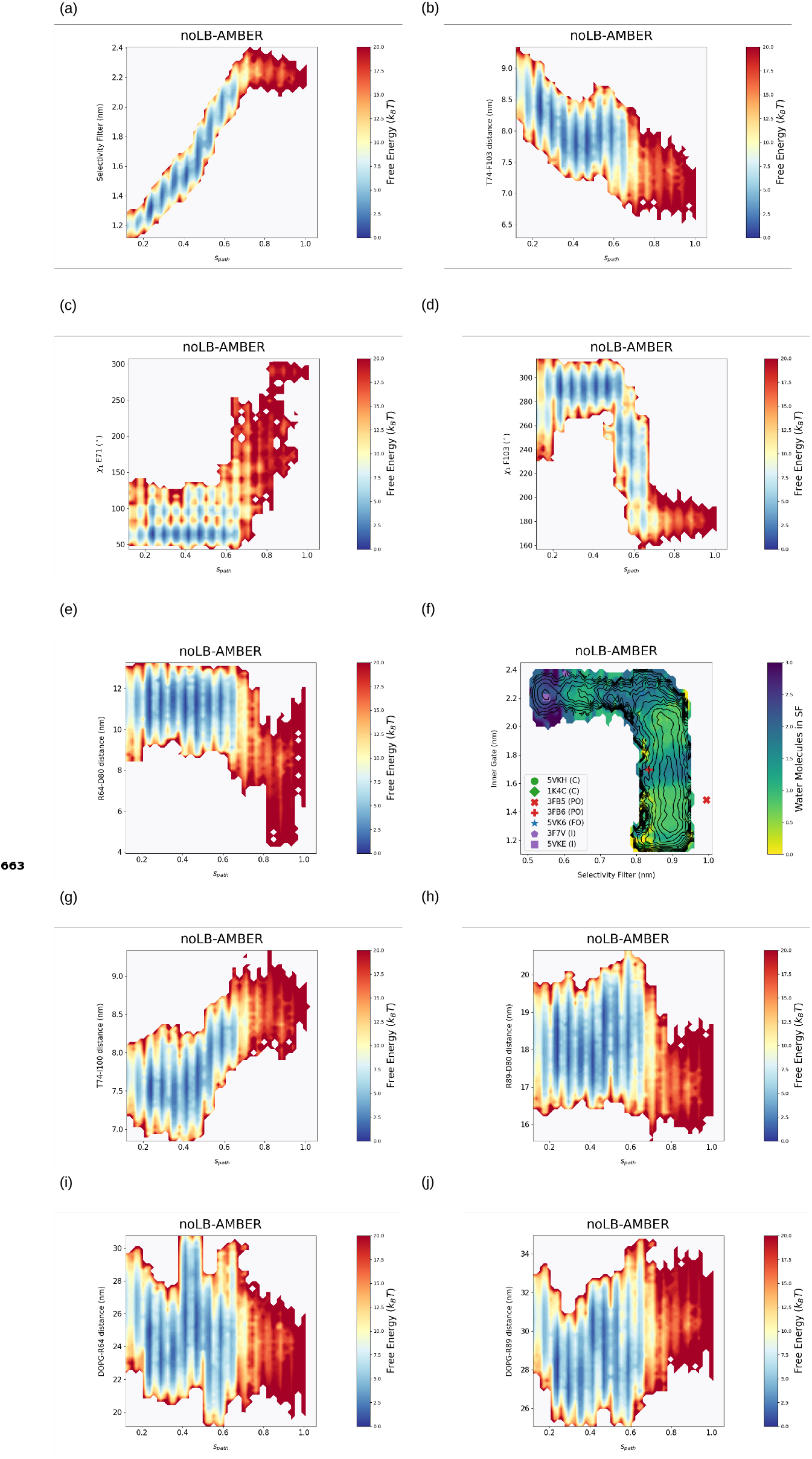
Reweighted free energy surfaces calculated from the string method with swarms of trajectories of the AMBER without lipids bound system. 2D free energy surfaces projected on the path cv (𝑠_𝑝𝑎𝑡ℎ_) and an additional CV of interest. These CVs of interest are: (a) Inner gate distance, (b) T74 CA - F103 CA distance, (c) E71 𝜒_1_ Janin angle, (d) F103 𝜒_1_ Janin angle, (e) R64 CZ - D80 CG distance, (g) T74 CA - I100 CA distance and (h) R89 CZ - D80 CG distance. (f) Projected weighted average number of water molecules inside the selectivity filter (sites S1-S4) on the free energy surface of the AMBER without lipid bound system (i) Copurifying DOPG P - R64 CZ distance (j) Copurifying DOPG P - R89 CZ distance.

**Figure 3—figure supplement 4.**
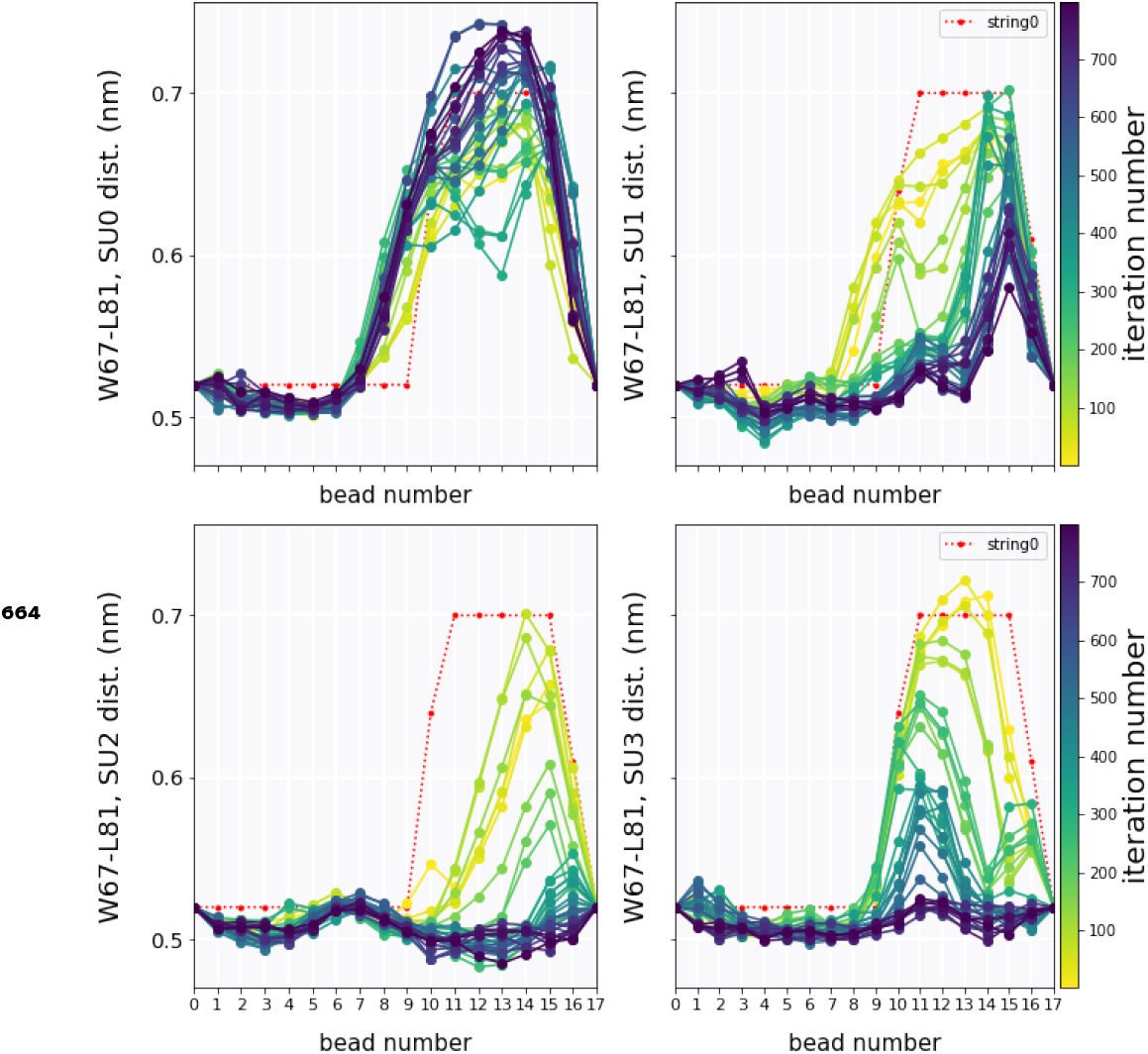
W67-L81 distance strings as a function of the string method iteration number for the AMBER system and with each graph representing the data for a particular subunit of the tetramer.

**Figure 3—figure supplement 5.**
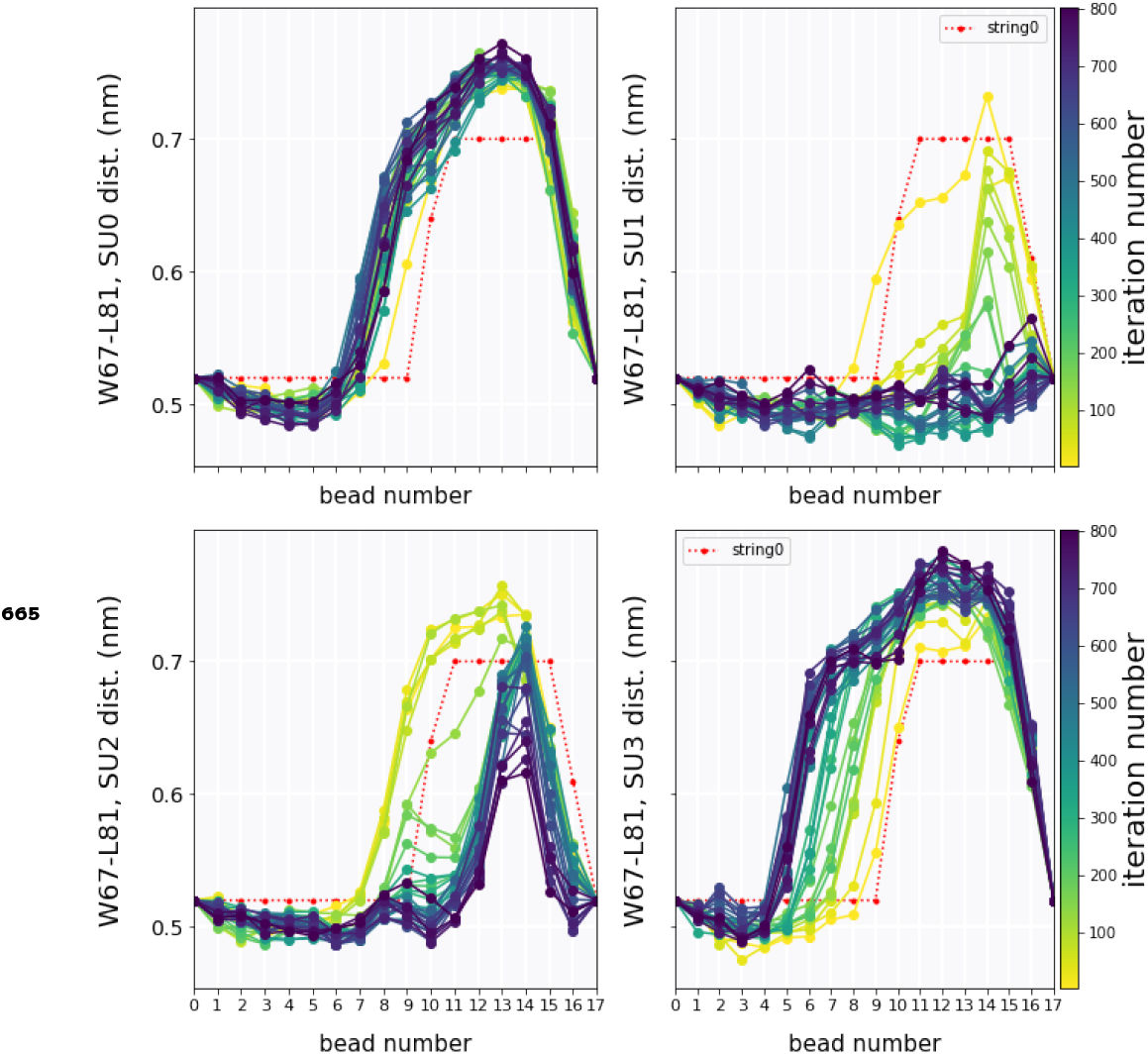
W67-L81 distance strings as a function of the string method iteration number for the CHARMM system and with each graph representing the data for a particular subunit of the tetramer.

**Figure 3—figure supplement 6.**
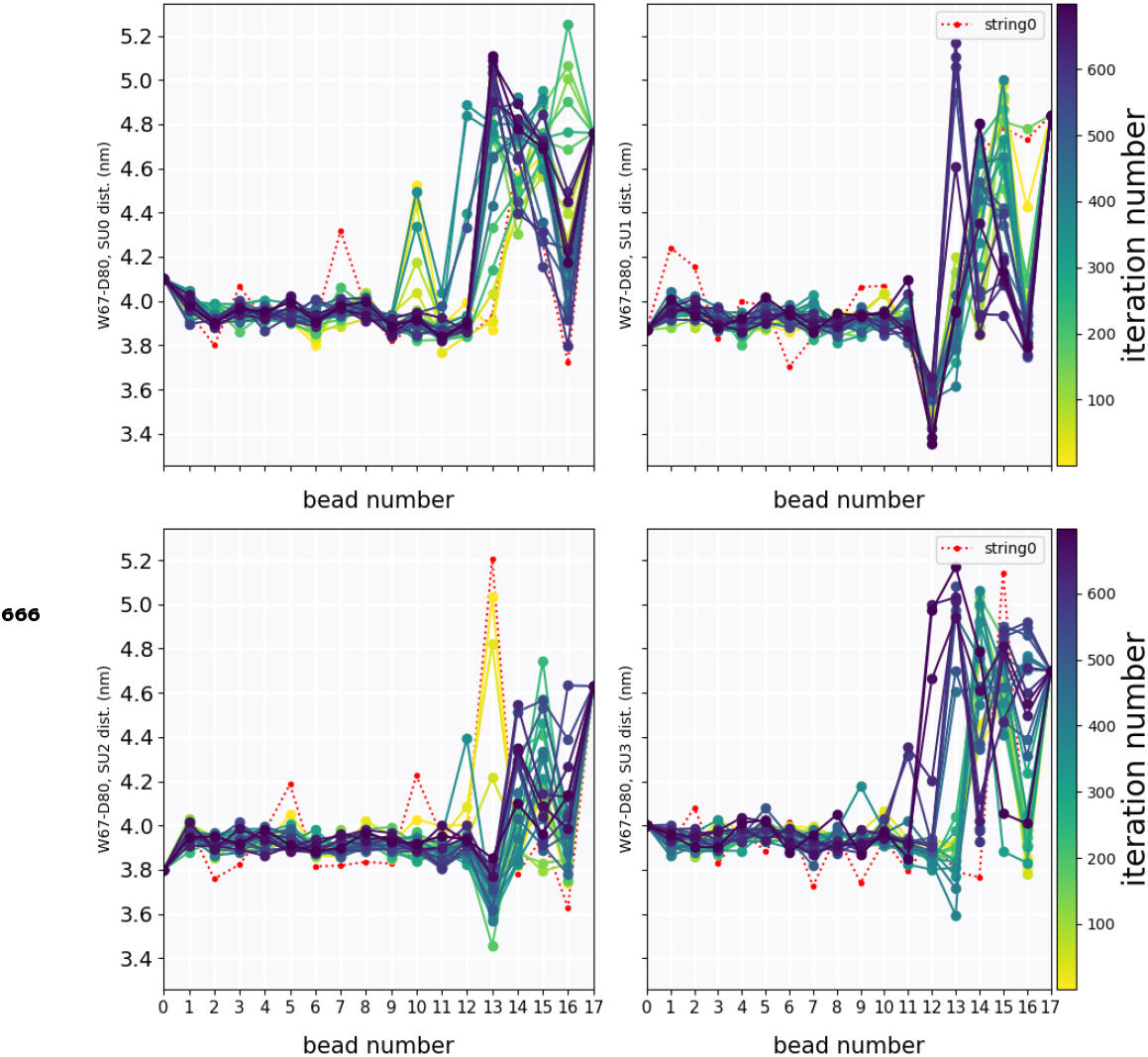
W67-D80 distance strings as a function of the string method iteration number for the AMBER system and with each graph representing the data for a particular subunit of the tetramer.

**Figure 3—figure supplement 7.**
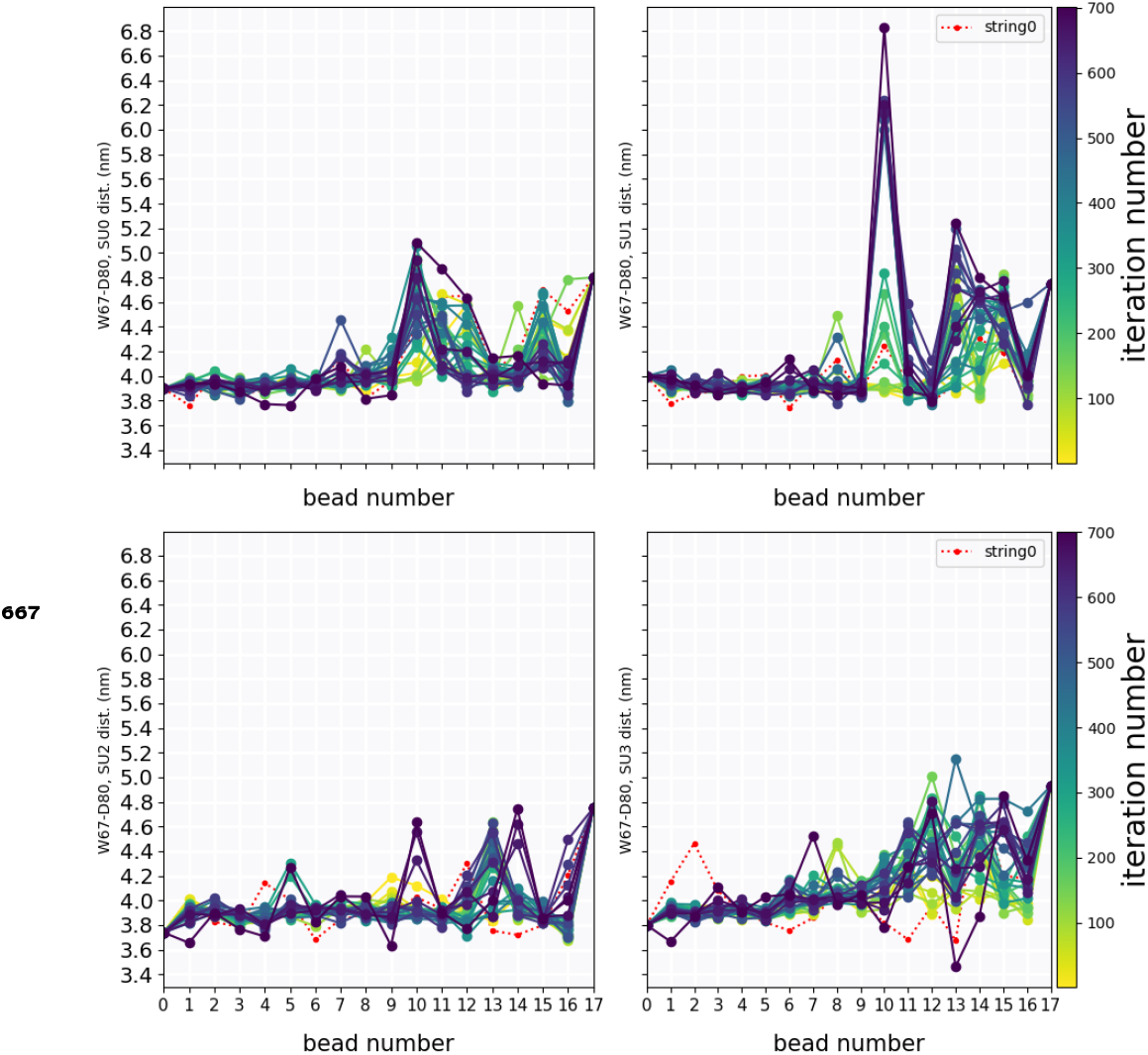
W67-D80 distance strings as a function of the string method iteration number for the CHARMM system and with each graph representing the data for a particular subunit of the tetramer.

**Figure 3—figure supplement 8.**
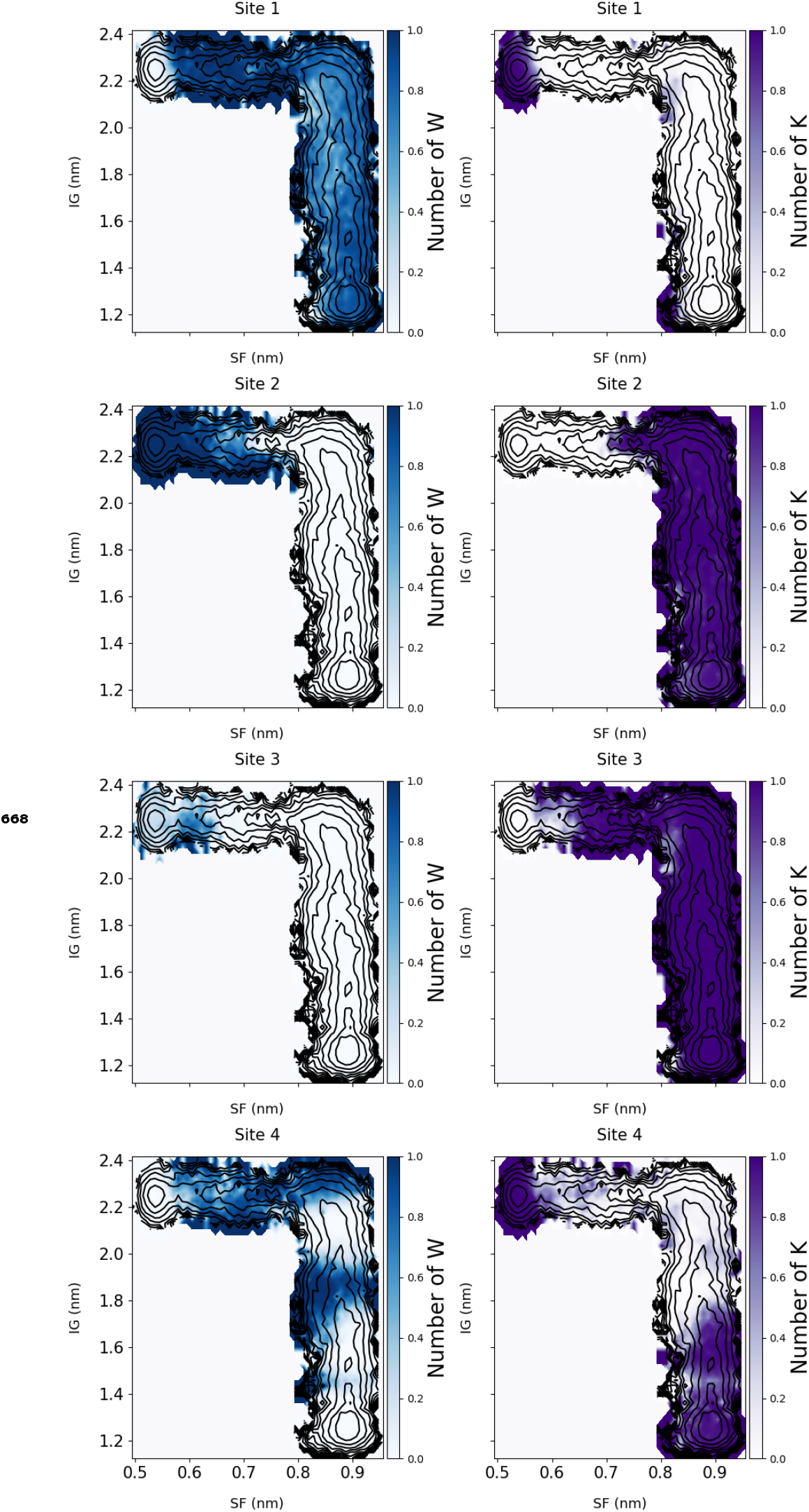
Projected weighted average number of water molecules (blue) or potassium ions (purple) inside a particular selectivity filter sites, S1 to S4 (rows), on the IG vs SF free energy surface of the AMBER with lipid bound system.

**Figure 3—figure supplement 9.**
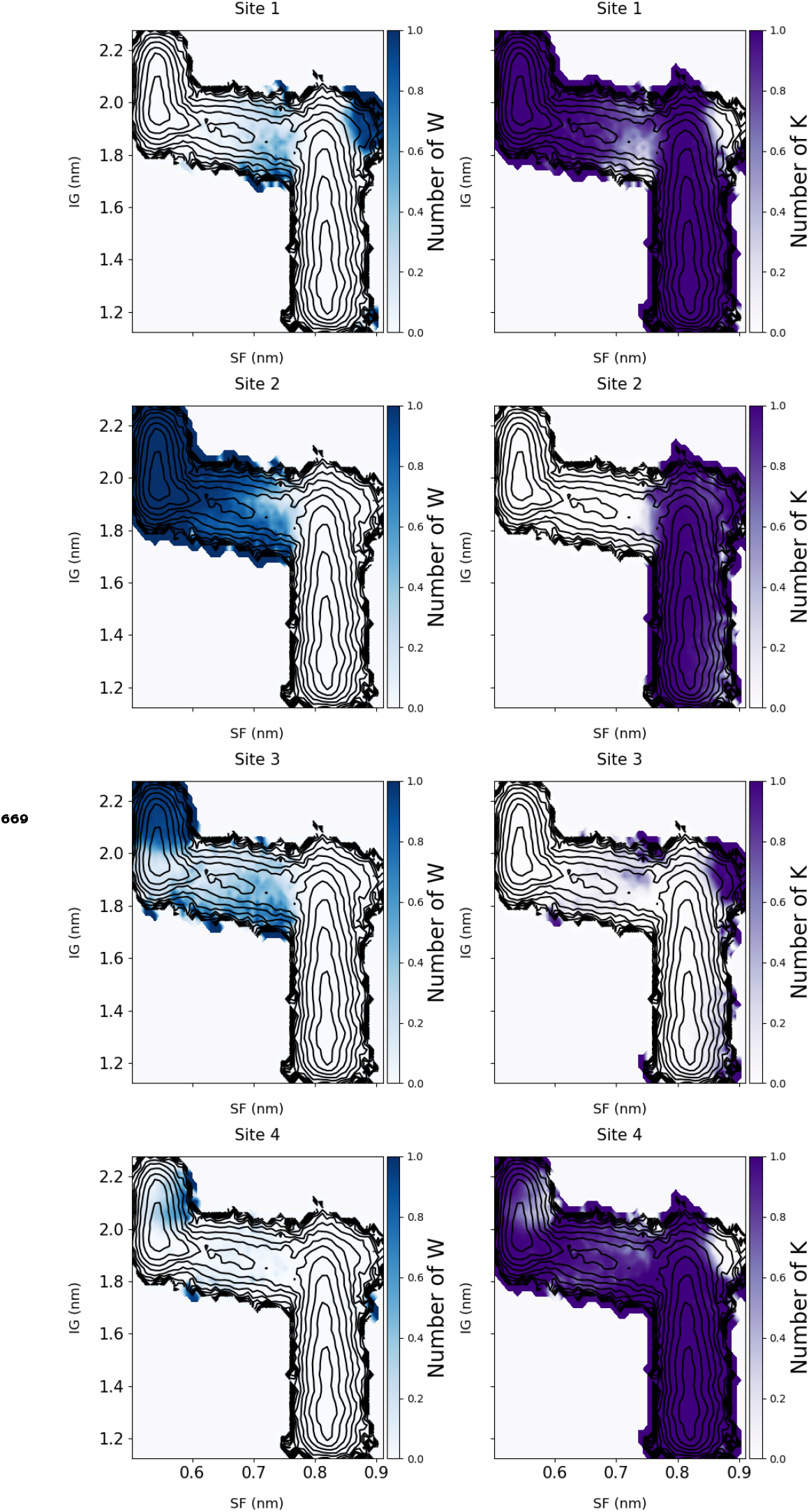
Projected weighted average number of water molecules (blue) or potassium ions (purple) inside a particular selectivity filter sites, S1 to S4 (rows), on the IG vs SF free energy surface of the CHARMM with lipid bound system.

**Figure 3—figure supplement 10.**
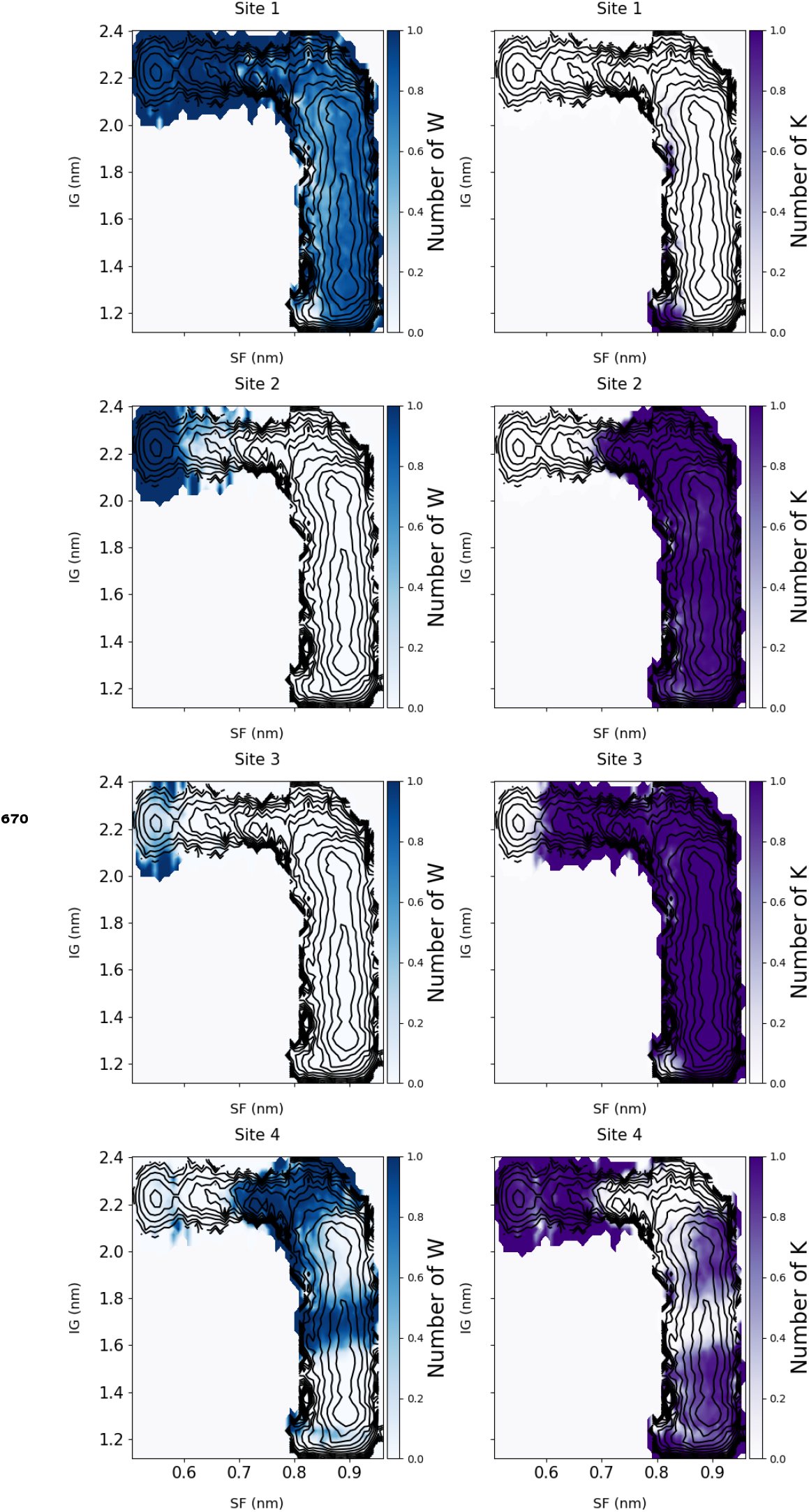
Projected weighted average number of water molecules (blue) or potassium ions (purple) inside a particular selectivity filter sites, S1 to S4 (rows), on the IG vs SF free energy surface of the AMBER without lipid bound system.

## Notes

https://github.com/delemottelab/KcsA\_string\_method\_FES

https://osf.io/snwbc

### Competing Interest Statement

The authors have declared no competing interest.

